# Regulation of Arabidopsis polyamine acetylation by NATA1 and NATA2

**DOI:** 10.1101/2024.03.04.583282

**Authors:** Umar F. Shahul Hameed, Yann-Ru Lou, Jian Yan, Francisco Javier Guzman Vega, Ekaterina Aleksenko, Pierre Briozzo, Solange Morera, Georg Jander, Stefan T. Arold

## Abstract

Polyamines have vital functions in organisms, including bacteria, plants, and animals, with key roles in growth, development, and stress responses. Spermine/spermidine *N*1-acetyl transferases (SSATs) regulate polyamine abundance by catalysing their *N*-acetylation, thereby reducing the pool of polyamines and producing other bioactive components. The regulatory mechanisms controlling SSAT enzymes are incompletely understood. Here, we investigate the biological role and regulation of the two SSAT isoforms present in *Arabidopsis thaliana*, *N*-ACETYLTRANSFERASE ACTIVITY (NATA) 1 and 2. We show that NATA2 is a heat-stable isoform, induced in response to heat. Intriguingly, a *nata2* knockout mutation proved beneficial for growth and pathogen defence under heat stress in Arabidopsis, aligning with the stress-mitigating effect of polyamines. In contrast, the double knockout of *nata1* and *nata2* was lethal, highlighting the essential role of basal SSAT activity. Our numerous crystal structures of both NATAs, supported by functional assays, revealed that stress-produced acidic metabolites can selectively inhibit polyamine acetylation by occupying the NATA substrate-binding pocket. This environment-responsive regulation mechanism may allow Arabidopsis to adjust the deleterious action of NATAs under stress conditions, without eliminating the enzyme. More generally, metabolite-ensemble inhibition may be a novel paradigm for non-genetic feedback regulation of plant enzymes.

## INTRODUCTION

Polyamines are abundant nitrogen-containing metabolites that mediate a wide variety of physiological functions in eukaryotes, bacteria, and archaea. In higher plants, the most common polyamines are the diamines putrescine and cadaverine, the triamine spermidine, and the tetraamine spermine (Kim et al., 2014; Regla-Márquez et al., 2016; Nahar et al., 2016; Sobieszczuk-Nowicka, 2017; Takahashi et al., 2018) (**Figure 1A**). These compounds play a central role in many aspects of plant physiology by influencing transcription, RNA modification, protein synthesis, enzyme activity, and signalling pathways (Evans and Malmberg, 1989; Galston et al., 1997; Kusano et al., 2007; Kusano et al., 2008; Takahashi and Kakehi, 2010; Wimalasekera et al., 2011; Sagor et al., 2015). In most plants, putrescine, the precursor for the synthesis of higher-order polyamines, is synthesised from arginine in two parallel pathways involving arginine decarboxylase, or *via* the non-proteinogenic amino acid L-ornithine (hereafter referred to as ornithine for simplicity), through the ornithine decarboxylase (**Figure 1A**) (Galston and Sawhney, 1990; Bagni and Tassoni, 2001). However, ornithine decarboxylase is absent in the model plant *Arabidopsis thaliana* (Arabidopsis) (Hanfrey et al., 2001).

**Figure 1.**
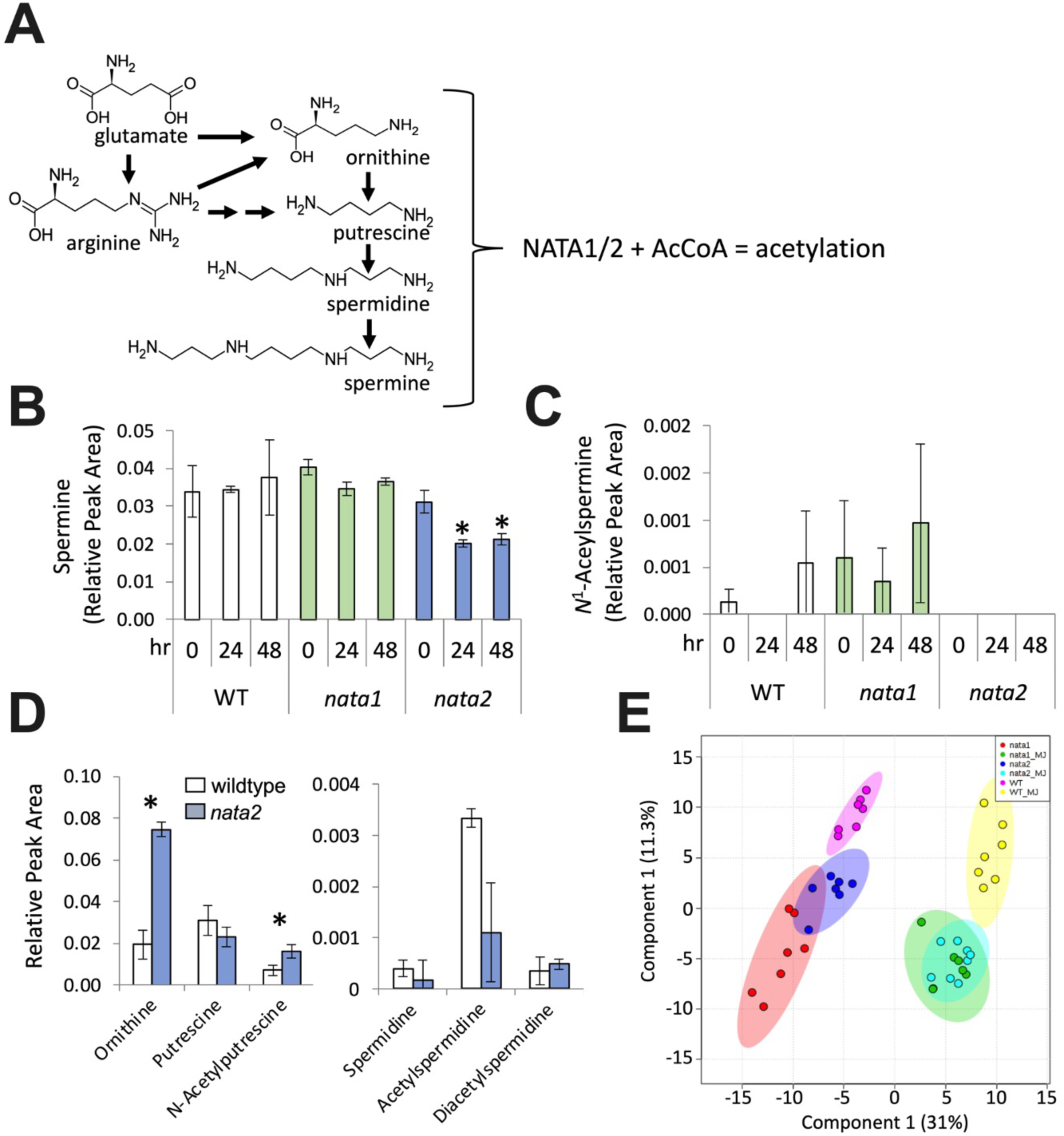
*In vivo* effects of *nata1* and *nata2* mutations. **A**) Pathways for biosynthesis of ornithine, putrescine, spermidine, and spermine, the targets of acetylation by NATA1 and NATA2. **B**) Spermine in content of wild-type Col-0, *nata1*, and *nata2* Arabidopsis seeds imbibed for 0, 24, and 48 hr, mean +/- s.e. of N = 3, *P < 0.05 relative 0 hour time point, Mann-Whitney U-test. **C**) *N^1^-acetylspermidine* concentration of Arabidopsis seeds imbibed for 0, 24, and 48 hr, mean +/- s.e. of N = 3. **D**) Metabolite concentrations in roots of wild-type Col-0 and *nata2* mutant Arabidopsis. Mean =/- s.e. of N = 3, *P < 0.05, *t*-test. **E**) Partial least squared discriminant analysis (PLS-DA) of HPLC-MS metabolite profiling data of wild-type Col-0 (WT), *nata1,* and *nata2* rosette leaves, untreated controls (salmon, blue, and pink) and four days after methyl jasmonate (MJ) treatment (green, pale blue, yellow). Ovals indicate 95% confidence regions, as determined by Metaboanalyst 3.0.

Polyamine abundance is regulated through import, export, synthesis, degradation, and chemical modification (Rhee et al., 2007; Zahedi et al., 2022). Spermidine/spermine *N*^1^-acetyltransferases (SSATs) are GCN5-related N-acetyltransferases (GNATs) that use acetyl-coenzyme A (acCoA) as a cofactor to transfer an acetyl to the aminopropyl moieties of polyamines, thus reducing the active pool of polyamines. SSATs have been identified in all kingdoms, including bacteria, protozoans, animals, and plants (Pegg, 2008). Polyamine acetylation can also create compounds with new functions and, despite being widely viewed as accumulative products, acetylated polyamines can influence plant metabolism by acting as pathway intermediates that facilitate turnover and translocation of polyamines (Tavladoraki et al., 2012; Moschou et al., 2012). Acetylated polyamines, suggestive of SSAT activity, have been identified in several plant species (Mesnard et al., 2000; Bagni and Tassoni, 2001; Fliniaux, 2004; Hennion et al., 2012; Tavladoraki et al., 2012; Lou et al., 2016). In Arabidopsis, *N*^1^-acetylspermidine was reported to be as abundant as spermidine in both roots and above-ground tissue (Kamada-Nobusada et al., 2008).

Three-dimensional (3D) structures of SSATs from bacteria, animals and plants have been deposited in the Protein Data Bank (PDB), but analyses have only been published for those from mouse, human, and the moss *Physcomitrium patens* (**Supplementary Table 1**) (Bewley et al., 2006; Han et al., 2006; Hegde et al., 2007; Montemayor and Hoffman, 2008; Bělíček et al., 2023). All SSATs consist of a central β-sheet flanked by helices, and form an intertwined obligate dimer through large surface contacts and arm exchange of the *N*-terminal β-strand. Differences between SSAT structures in the free form and bound to cofactors and/or substrates (**Supplementary Table 1**) suggest that catalysis involves conformational changes. So far, these structural transitions have not yet been described for the same protein, precluding a precise analysis of structural transitions and their role in catalysis.

Plant SSATs distinguish themselves from animal and bacterial enzymes by a 35-amino acid insert (**Supplementary Figure 1A**). In the sole SSAT 3D structure reported with this insert, from *P. patens* (Bělíček et al., 2023), it forms an additional β-strand and a loop, with the latter being partially disordered in the crystal structure (shown red in **Figure 4A, Supplementary Figure 1B**). The proximity of the additional β-strand to the cofactor binding site suggests that the insert affects catalysis (Bělíček et al., 2023) but the nature of the effect remains unclear. Indeed, the presence of the plant insert does not correlate with the catalytic rate or substrate preference of the enzyme (Bělíček et al., 2023).

SSATs generally display low turnover rates and a broad substrate profile, ranging from diamines to spermine, also including 1,3-diaminopropane and the non-proteinogenic amino acids ornithine and thialysine (Adio et al., 2011; Jammes et al., 2014; Lou et al., 2016; Mattioli et al., 2022). Nonetheless, several studies show different substrate preferences for some SSATs. These differences in substrate preference were apparent not only between kingdoms but also between species within kingdoms. For example, the plant SSATs from moss, maize and Arabidopsis were shown to acetylate a range of polyamines, including 1,3-diaminopropane, putrescine, spermidine and spermine, as well as the precursor amino acid ornithine (Bělíček et al., 2023). However, the substrate preferences differed significantly between plant species, among different parts of the same plant, and depending on whether plants were subjected to osmotic stress or water stress (Sen et al., 2018). Moreover, differences were also observed among paralogues from the same species. Human SSAT2, for example, was reported to acetylate thialysine rather than spermine or spermidine, whereas both polyamines were good substrates for human SSAT1 (Della Ragione and Pegg, 1983; Coleman et al., 2004; Hegde et al., 2007; Pegg, 2008). As an explanation, it was suggested that the polyamine processing profile is dependent on small changes in the protein active site and surrounding areas, allowing polyamine acetylation to be species-specific (Mattioli et al., 2022). Yet, even for the same SSAT, different researchers observed markedly different substrate preferences, suggesting experimental factors can modulate catalysis (e.g. buffer, pH or temperature). For instance, the best substrate for the Arabidopsis SSAT *N*-ACETYLTRANSFERASE ACTIVITY1 (NATA1; At2g39030) was reported to be either ornithine, thialysine, or putrescine (Adio et al., 2011; Jammes et al., 2014; Lou et al., 2016).

Although it is expected that the activity of SSATs is tightly controlled, the basis for this regulation remains poorly defined and controversial. In *P. patens*, SSAT is not significantly upregulated under various abiotic stresses (Bělíček et al., 2023), whereas the Arabidopsis *NATA1* expression is upregulated by a variety of stresses, including cold, salinity, drought, wounding, as well as addition of jasmonate, cytokinins, and polyamines (Adio et al., 2011; Jammes et al., 2014; Lou et al., 2016; Mattioli et al., 2022).

A gene duplication event in Arabidopsis and closely related species led to the presence of two *NATA* genes, which encode proteins with approximately 80% amino acid sequence identity, (Lou et al., 2016). Unlike *NATA1*, *NATA2* (At2g39020; likely the evolutionarily ancestral gene), was not upregulated by cytokinins and polyamines (Lou et al., 2016; Mattioli et al., 2022). The selective advantage of the *NATA* gene duplication in Arabidopsis, and the biological role of NATA2 remain unclear. Indeed, and surprisingly, the deletion of *NATA1* increases both growth and pathogen resistance of Arabidopsis (Lou et al., 2016), in line with the importance of polyamines for growth and stress resistance. The upregulation of *NATA1* in Arabidopsis under conditions where SSAT activity is unfavourable, and the presence of a second highly similar gene, are intriguing. Moreover, considering that the catalytic activity of NATA1 and NATA2 on polyamines is generally relatively low, and that there are also other pathways for polyamine catabolism (through copper-containing amine oxidases (CuAOs) and flavin-containing polyamine oxidases (polyamineOs) (Tavladoraki et al., 2016), it is unclear how important NATA1 and NATA2 are for plant growth.

Here, we investigated the biological role of NATA2 in Arabidopsis and explored factors regulating the activity of both NATA paralogues. We found that NATA2 exhibited greater heat stability than NATA1 and was upregulated under heat stress. Surprisingly, the absence of the *NATA2* gene enhanced growth and pathogen resistance of Arabidopsis at high temperatures. This phenomenon mirrored the increased expression of NATA1 induced by the plant defence signalling molecule jasmonic acid, even though microbial growth was reduced in *nata1* mutant Arabidopsis. Notably, our experiments highlighted that the deletion of both *NATA* genes is lethal, underscoring their vital housekeeping functions. Structural and biochemical analyses provided a detailed description of the NATA1/2 catalytic cycle. Furthermore, these investigations suggested that the NATA active site has evolved to permit catalytic inhibition by a set of acidic metabolites, prevalent in plants under stress conditions. Overall, our results elucidate the role and regulation of Arabidopsis SSATs, and propose a general mechanism for metabolite sensing that allows selective attenuation of SSAT activity under biotic and abiotic stresses.

## RESULTS

### Knockout of nata2 changes the Arabidopsis metabolite profile

We characterised the catalytic function of NATA2 *in planta* using an Agrobacterium T-DNA insertion knockout of At2g39020 (Salk_092319; *nata2-1*), which has been described previously (Adio et al., 2011; Lou et al., 2016). The mutant showed no obvious defects and initiated flowering at the same time as the wild-type Columbia-0 (Col-0) under our growth conditions. We did not observe significant differences in the ornithine, putrescine, or acetylputrescine concentrations when comparing rosette leaf extracts of *nata2-1* and wild-type plants (**Supplementary Figure 2A**). Under these conditions, spermine and spermidine were not abundant enough in Arabidopsis leaves for reliable quantification. The absence of a polyamine phenotype might be due to the low expression level of NATA2 in rosette leaves, especially under non-stressed conditions (Waese et al., 2017).

The relative expression of *NATA2* was highest in imbibed Arabidopsis seeds (Waese et al., 2017). Therefore, we used HPLC-MS to analyse polyamine content in imbibed seeds of wild-type Columbia-0 (Col-0) Arabidopsis, the *NATA2* knockout line (*nata2-1*), and a *NATA1* knockout line (GK-256F07; *nata1-1*). There was a small but significant decrease in spermine content in germinating *nata2-1* seeds after 24 and 48 hours (**Figure 1B**). Whereas wild type and *nata1-1* Arabidopsis seeds generally showed low levels of acetylspermine (close to the detection limit of our assay), this compound was not detected in *nata2-1* seeds 0, 24, or 48 hours after imbibition (**Figure 1C**).

To determine NATA2 effects in roots, where spermine and acetylspermine were reported to be equally abundant (Kamada-Nobusada et al., 2008), we extracted root tissue from *nata2-1* and wild-type Arabidopsis for analysis. Ornithine and acetylputrescine were more abundant in *nata2-1* roots than in wild-type roots (**Figure 1D**). There were no significant changes in the abundance of putrescine, spermidine, acetylspermidine, or diacetylspermidine, and neither spermine nor acetylspermine were detected in this assay. We concluded that the absence of NATA2 had generally mild or no effects under the conditions tested.

### The double knockout of nata1 nata2 is lethal, showing that NATA function is essential

The relatively mild phenotype of individual *nata1* and *nata2* knockouts, and the 81 % amino acid sequence similarity between NATA1 and NATA2, suggested a significant functional redundancy between these orthologues. Indeed, partial least-squares discriminant analysis (PLS-DA) of non-targeted LC-TOF-MS metabolite profiling data from rosette leaves showed that the metabolite profiles of *nata1* and *nata2* are more similar to each other than to wild-type Col-0, both in uninduced plants and after methyl jasmonate elicitation (**Figure 1E**; **Supplementary Data Set 1**).

Given the similar metabolic effects of *nata1* and *nata2* mutations, we attempted to produce a homozygous double knockout mutant. As *NATA1* and *NATA2* are directly adjacent to one another on Arabidopsis chromosome 2, the expected low level of meiotic recombination between mutant alleles precluded the use of traditional crossing techniques. Transcription Activator-Like Effector Nucleases (TALENs; (Cermak et al., 2011)) were previously used to create a *NATA2* knockout mutation in the *nata1-1* mutant background (Christian et al., 2010; Christian et al., 2013). We confirmed the presence of a heterozygous, heritable 7 bp deletion in the NATA2 coding region (*nata2-2*) in a homozygous *nata1* mutant background (**Supplementary Figure 2B**). However, despite multiple generations of screening, we did not obtain any homozygous *nata1-1 nata2-2* double mutants from selfed *nata1-1*/*nata1-1* NATA2/*nata2-2* plants. Although developing seeds and seed pods looked normal and seed pods were not missing seeds, suggesting that the double mutation is not embryonic lethal, we observed a reduced germination rate in seeds from two heterozygous *nata1-1*/*nata1-1 NATA2*/*nata2-2* lines. In one case 147 out of 190 seeds (77%) successfully germinated and in the other 81 out of 117 (69%) germinated. This is similar to the expected 3:1 live:dead ratio if the homozygous *nata1-1 nata2-2* double mutation is lethal during germination or early in development (P < 0.05, chi-squared test). PCR-based genotyping of plants that had germinated showed no homozygous *nata1-1*/*nata1-1 nata2-2*/*nata2-2* double mutants. Instead, successfully germinated plants had the other two expected genotypes from this cross, either *nata1-1*/*nata1-1 NATA2*/*nata2-2* or *nata1-1*/*nata1-1 NATA2*/*NATA2*, and had no visible defects compared to wild-type Arabidopsis.

### The nata2-1 mutant is less sensitive to heat stress during hypocotyl elongation

Despite a relatively high expression level in imbibed seeds and young seedlings (**Supplementary Figure 2C**) (Waese et al., 2017), the role of NATA2 in seedling development is largely unknown. Under 23°C growth conditions, Arabidopsis *nata2-1* seedlings showed no obvious growth phenotype compared to wild type seedlings (**Supplementary Figure 2D**). However, because *NATA2* expression is heat-inducible (**Supplementary Figure 3**) (Schmid et al., 2005; Kilian et al., 2007), we hypothesised that NATA2 plays a role in regulating heat stress responses during seed development by fine-tuning polyamine levels.

To test this hypothesis, we performed a hypocotyl elongation experiment similar to the one reported by (Shen et al., 2016). No hypocotyl length differences between mutant and wild type were observed when seedlings were grown at 23°C (**Figure 2A**). By contrast, the slightly longer hypocotyls of *nata2-1* seedlings compared to wild-type Col-0 at 37°C suggest that the mutants are more tolerant of heat stress. To determine whether the observed phenotype can be the result of altered polyamine titer due to NATA2 activity, we analysed the polyamine profile of *nata2-1* mutant and wild type seedlings after heat stress. Although wild-type seedlings accumulated more spermine than *nata2-1* mutants under heat stress (**Figure 2B**), the abundance of *N*1-acetylspermine was too low to be reliably detected. There were no significant differences in the abundance of seedling ornithine, putrescine, acetyl-putrescine, or spermidine in this experiment.

**Figure 2.**
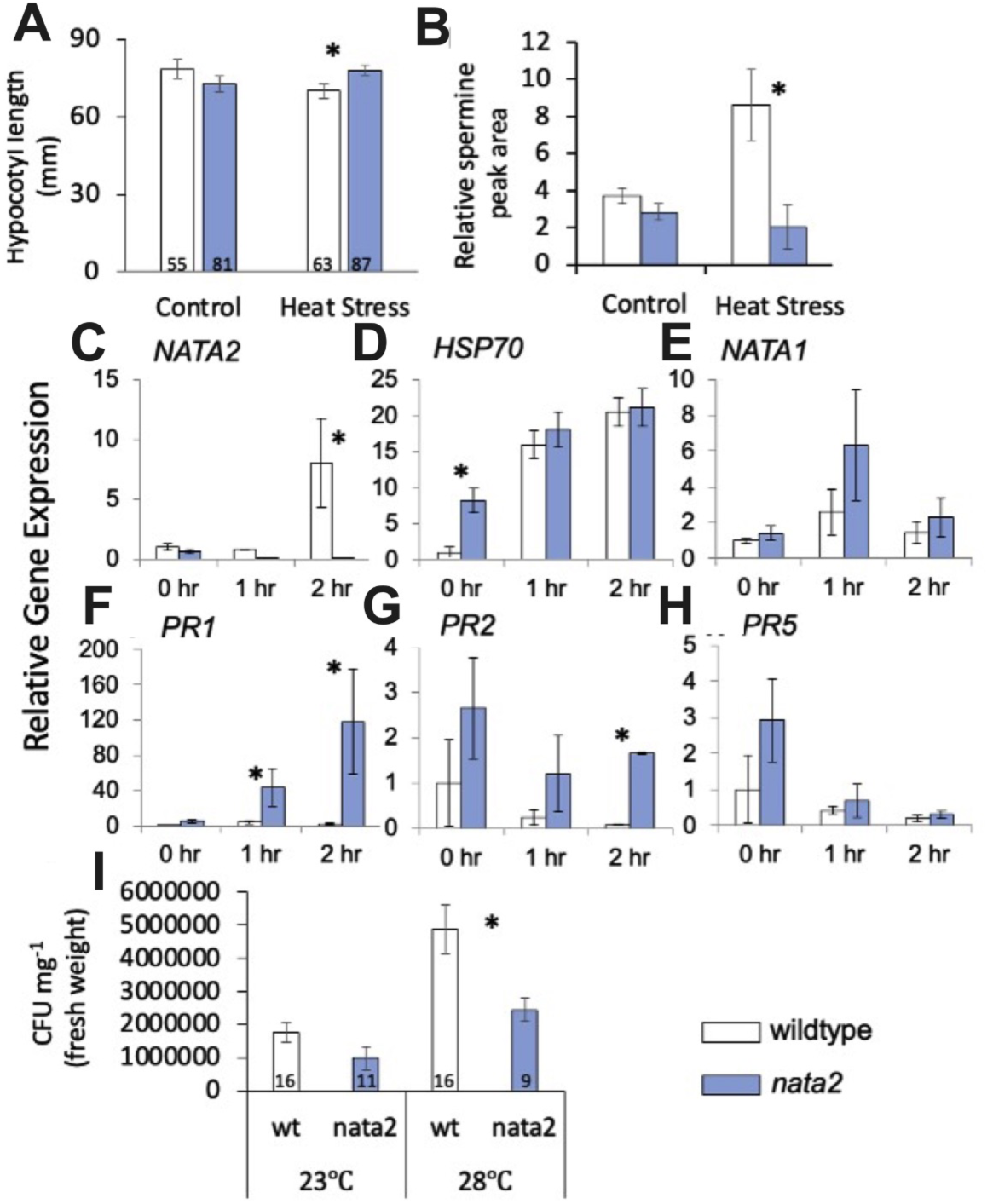
Altered heat stress and defence responses in *nata2* mutants. **A**) Hypocotyl elongation in control seedlings at 23°C and heat-stressed seedlings at 37°C . Mean +/- s.e., numbers in bars indicate sample sizes, *P < 0.05, two tailed *t-*test. **B**) Spermine accumulation in 7-day-old wild type and *nata2* mutant seedlings after three hours of 37°C heat stress. Mean ± SE of N = 4-5, *P < 0.05, two tailed *t-*test. Transcript abundance of (**C**) *NATA2*, (**D**) *HSP70*, (**E**) *NATA1*, (**F**) *PR1*, (**G**) *PR2*, and (**H**) *PR5* in wild type and *nata2* mutant seedlings, 0, 1, and 2 hours after heat stress at 37°C. Gene expression was measured by quantitative RT-PCR. Mean ± SE of N = 6-12. *P < 0.05, two tailed *t-*test. **I**) Bacterial titers in wild-type Col-0 and *nata2* rosette leaves four days after infiltrating 10^5^ colony forming units (CFU) ml^-1^ culture of *Pseudomonas syringae* strain DC3000. Mean +/- s.e., numbers in bars indicate sample sizes, *P < 0.05, two tailed *t*-test.

To further investigate the *nata2* mutant effects in Arabidopsis seedlings, quantitative RT-PCR assays were used to determine whether known stress-responsive genes are differentially regulated compared to Col-0. Consistent with previously reported results (Waese et al., 2017), *NATA2* expression increased within two hours after the application of 37 °C heat stress (**Figure 2C**). Little or no *NATA2* transcripts were detected in the *nata2-1* mutant, confirming the knockout effects of the T-DNA insertion. Expression of *HSP70*, a commonly used marker for the heat stress responses in Arabidopsis, was not higher in the *nata2* mutant than in wild type after heat stress (**Figure 2D**), though it was elevated at t = 0 before the stress was applied. Both groups of seedlings also showed comparable *NATA1* transcript abundance (**Figure 2E**). Interestingly, the *nata2-1* mutation caused elevated expression of *PR1* and *PR2*, two commonly used markers for pathogen defence responses in Arabidopsis (**Figure 2F, G**). There was, however, no increase in the expression of another defence-related gene, PR5 (**Figure 2H**).

### NATA2 is detrimental for plant defence under heat stress

The observation of increased *PR1* and *PR2* expression in *NATA2*-deficient plants suggested elevated defence against biotrophic pathogens. This is consistent with prior observations that polyamine abundance in plant cells correlates with defence against pathogens (Yoda et al., 2006; Yoda et al., 2009; Kim et al., 2014; Rossi et al., 2014; Lou et al., 2016). Hence, we hypothesised that, given the regulatory function of polyamine homeostasis, NATA2 plays a role in the crosstalk between abiotic stress and biotic stress response pathways, *i.e.*, between heat stress and pathogen defences. To test the hypothesis, 18-day-old wild type and *nata2* mutant Arabidopsis were infected with *P. syringae* at 23°C and 28°C, a temperature that is suboptimal for normal *A. thaliana* growth. Four days after infiltration, bacterial counts were similar at 23°C but were higher in wild-type Col-0 than in *nata2-1* plants at 28°C (**Figure 2I**).

### NATA1 and NATA2 have a similar broad substrate preference in vitro

To complement the *in vivo* results, we produced *A. thaliana* NATA1 and NATA2 proteins in *E. coli* to investigate enzyme functions under controlled *in vitro* conditions. We produced full-length NATA2, and NATA1 and NATA2 constructs where we deleted the flexible N-terminal extension (residues 1-18 and 1-29, respectively), that we named NATA1Δ and NATA2Δ. Almost all *in vitro* analyses were based on NATA1Δ and NATA2Δ constructs, except for one crystal structure, which was based on full-length NATA2. NATA1Δ showed highest relative catalytic activity on thialysine, 1,3-diaminopropane, and spermine, which was narrowly better than its activity on ornithine, putrescine, and lysine (**Figure 3A, Table 1, 2**). When tested in 50 mM Tris–HCl pH 8.5, 150 mM NaCl (referred to as buffer Tris8.5) at 25 °C, NATA2Δ showed the same relative activity profile as NATA1Δ on all substrates except for ornithine, which it acetylated at very low levels. This broad substrate preference was not specific to the buffer used, as we obtained a similar broad acetylation profile, including NATA2Δ’s poor catalysis of ornithine, under different conditions (100 mM Tris-HCl, pH 7.5, 2 mM EDTA and 10% glycerol, referred to as Tris7.5, at 30 °C) (**Figure 3B**). However, the overall catalytic efficacies were reduced in Tris7.5, consistent with earlier observations that NATA2Δ activity is higher at a pH range of 8.5 – 9.5 (**Supplementary Figure 4A**) (Mattioli et al., 2022).

**Figure 3.**
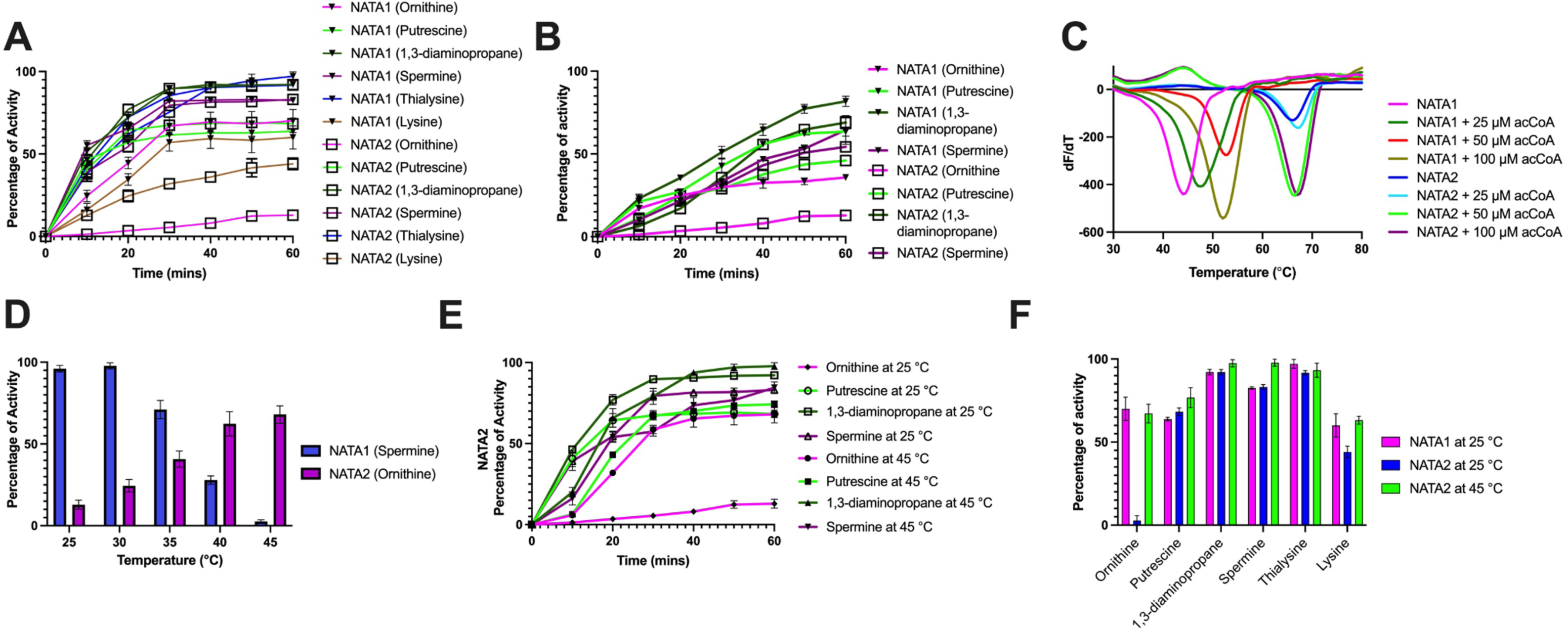
NATA1Δ and NATA2Δ substrate preference and heat stability. Catalytic activity of NATA1Δ and NATA2Δ on their substrates were measured (**A**) in buffer Tris8.5 at 25°C and (**B**) in buffer Tris7.5 at 25°C. Concentrations of substrate and acCoA were 5 mM and 0.5 mM, respectively. Protein concentrations were 3 µM. **C**) Stability of NATA1Δ and NATA2Δ assayed by DSF in the presence of different concentrations of acCoA, measured in buffer Tris8.5. **D**) Catalytic activity of NATA1Δ and NATA2Δ on spermine or ornithine at 25 and 45°C in buffer Tris8.5. Concentrations used as in (A). **E**) Catalytic activity of NATA2Δ on its substrates at 25 °C and 45 °C in buffer Tris8.5. Concentrations as in (A). **F**) Endpoint histogramme, based on data from (E).

**Table 1.** Relative activity of NATA1Δ and NATA2Δ against substrates. Measurements were carried out in Tris8.5 buffer at 25 and 45 °C

| Relative activity (%) |  |  |  |
| --- | --- | --- | --- |
| Substrate | NATA1Δ (25 °C) | NATA2Δ (25 °C) | NATA2Δ (45 °C) |
| Thialysine | 97.12 ± 2.67 | 91.79 ± 1.21 | 93.25 ± 4.31 |
| 1,3-diamino propane | 92.12 ± 1.67 | 92.12 ± 1.61 | 97.45 ± 2.16 |
| Spermine | 82.60 ± 0.79 | 83.13 ± 1.52 | 97.80 ± 2.12 |
| Putrescine | 63.79 ± 1.07 | 68.28 ± 2.27 | 76.75 ± 5.99 |
| Ornithine | 70.04 ± 7.04 | 12.87 ± 2.79 | 67.20 ± 5.63 |
| Lysine | 60.04 ± 7.03 | 44.04 ± 3.50 | 63.15 ± 2.35 |

### NATA2 is the heat stable form of NATA1

Having observed that *NATA2* expression was induced by heat (**Figure 2C**), we used differential scanning fluorimetry (DSF) to assess the stability of the purified *E. coli*–produced proteins. With a melting temperature *Tm* of 42.5 ± 0.7°C, apo NATA1Δ was markedly less stable than apo NATA2Δ (*Tm* = 66.5 ± 0.5°C). The addition of 50 µM acetyl CoA (acCoA), but not substrates, increased the heat stability of NATA1Δ by 10°C, whereas no stabilisation was observed for NATA2Δ (**Figure 3C, Supplementary Table 3A**). The difference in stabilisation was not caused by differences in binding energy, as both enzymes bound acCoA with a dissociation constant, *Kd*, of approximately 10 µM and similar entropy-dominated thermodynamics (**Supplementary Figure 4B,C**). The presence of substrates did not lead to a significant increase in *Tm* for either enzyme (**Figure 3C, Supplementary Table 3A**). In agreement with its *Tm*, the ability of NATA1Δ to acetylate spermine declined at temperatures above 30 °C, with a total loss of activity at 45 °C. Conversely, the NATA2Δ activity on ornithine increased until 45°C (**Figure 3C**). As a result, the substrate preference profile of NATA2Δ at 45°C resembled that of NATA1Δ at 25°C, including ornithine (**Figure 3E,F**). Hence, at elevated temperature, NATA2 recovers the activity loss of NATA1.

### Identifying the molecular basis for heat stability

To investigate the molecular basis for the observed properties of NATA1 and NATA2, we determined a total of eight different crystal structures at resolutions between 1.1 and 2.0 Å, three for NATA1Δ, four for NATA2Δ, and one using full-length NATA2. (**Supplementary Table 1, 2**). In the case of NATA1Δ, we obtained a structure in the presence of acCoA, and structures with CoA and either a natural (ornithine) or artificial (HEPES) ligand. We obtained NATA2 structures in the apo state for both full-length and NATA2Δ. In full-length NATA2 only residues 20–29 of the N-terminal extension were visible. These residues were loosely associated with the cofactor binding site of a crystallographically related NATA2 molecule, as illustrated by their above-average B-factors, supporting that the residues before the NATA2Δ core are flexible. We also determined NATA2Δ structures bound to spermine, or to HEPES and acCoA. Additionally, we obtained a NATA2Δ structure bound to a disulphide-linked di-CoA molecule that was produced *in cristallo* based on the presence of 2 mM CoA.

The catalytic domains of NATA1 and NATA2 adopt the intertwined α/β dimer structure observed for the bacterial, animal and moss SSAT enzymes (**Figure 4A**). The 35-residue insertion specific to the plant enzymes forms an additional β-strand (residues 97-103 and 105-111 for NATA1 and NATA2, respectively) similarly to what was observed for *P. patens* SSAT (Bělíček et al., 2023). The remaining insert residues form a loop from which F98, F101 and I111 attach to residues F72, F78 and Y159 of the NATA1 core (residues F106, F109 and I119 binds back to the residues of F79, F85, and Y167 in NATA2) (**Figure 4A, Supplementary Figure 5A**). The structure of the unattached part of the loop varies between all the protein chains, and lacks continuous electron density in most monomers, suggesting flexibility in solution. In one of the two molecules of the asymmetric unit of the NATA2Δ structure bound to di-CoA, the additional β-strand region (residues 114-109) was moved by 5 - 7 Å, losing its secondary structure (**Supplementary Figure 5B**). This conformation likely resulted from the insertion of the C-terminal helix of a neighbouring molecule, suggesting that the structure of the plant-specific insert at this position may be susceptible to ligands.

**Figure 4.**
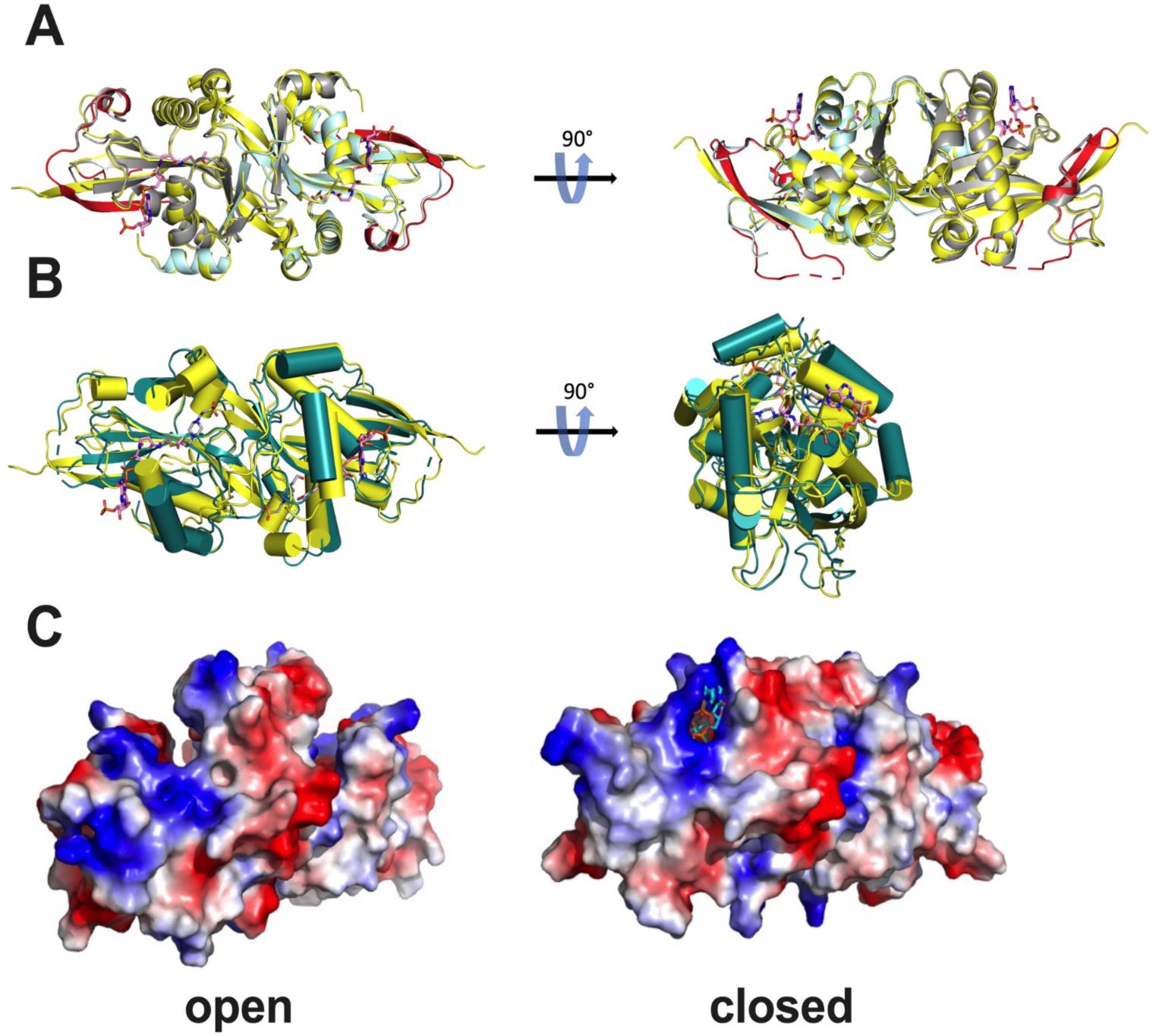
Structural conformations from NATA1Δ and NATA2Δ crystal structures. **A**) Superimposition of NATA1Δ (grey and cyan) and NATA2Δ (yellow) dimers. The plant-specific extension is shown in red. The CoA bound to NATA1Δ is shown as a stick model with carbons coloured in pink. The structure on the right is rotated 90⁰ along the x-axis. **B**) Superimposition of apo-NATA2Δ (dark green), adopting the open conformation, onto the closed structure of NATA2Δ (yellow) bound to HEPES and acCoA (both shown as stick model). The carbon atoms of acCoA and HEPES are coloured in pink and grey. Helices are shown as cylinders. Right figure shows the front view, as obtained by two orthogonal 90° rotations. **C**) Electrostatic surface of apo-NATA2Δ (left, open conformation) and NATA2Δ bound to HEPES and acCoA (right, closed conformation). Structures are roughly placed in the orientation of figure (A), right panel, but slightly rotated around the Y axis to better show the changes in the open and closed conformations.

In all structures that contain the cofactor, its α and β phosphates are coordinated in the same way by the backbone nitrates of the P-loop and the following helix (residues 161-166 and 169-174 for NATA1 and NATA2, respectively). The positions of the other parts of the cofactor vary according to the different structural contexts. Compared to other known SSAT structures, the P-loop of NATA1 and NATA2 contains an additional arginine (R161 and R169, respectively) which contributes to coordinating the 3’ phosphate in some structures. This arginine is present only in higher plants such as Arabidopsis and maize, but not in moss, human, or Pseudomonas SSATs, and may lead to variations in cofactor binding dynamics (**Supplementary Figure 5C**).

To investigate the molecular basis for the difference in thermal stability and stabilising effect of CoA, we searched for notable differences between the 76 % sequence-identical paralogues. The most prominent difference occurs in the centre of the dimer, close to the substrate binding site, where NATA1 residues V75, Q146, and V181 are substituted in NATA2 by larger and/or more hydrophobic residues (F82, P154, and Y189, respectively) (**Figure 4A, Supplementary Figure 6A**). Their importance in stabilising NATA2 is supported by stability calculations using FOLDX: Using the NATA2Δ apo structure as a template, the NATA2 sequence had a 23 kJ lower calculated energy than the NATA1 sequence. The *in silico* triple mutant increased or lowered the calculated energy for NATA1Δ and NATA2Δ, respectively, by more than 10 kJ (**Supplementary Figure 6B**). In agreement, the recombinantly produced F82V, P154Q, and Y189V triple mutation decreased the NATA2Δ *Tm* by 10 °C for the apo form, and increased the stabilising effect of acCoA significantly, akin to wild-type NATA1Δ. The NATA1Δ V75F/Q146P/V181Y mutant decreased the stabilising effect of acCoA (from 10°C to 6.5 °C) but failed to increase the *Tm* of the mutant protein (**Supplementary Figure 6C, Supplementary Table 3A**). We conclude that these three amino acid positions have a significant role in determining the protein stability but are insufficient to fully recapitulate the *Tm* differences.

### NATA1 and NATA2 crystal structures show open and closed conformations

NATA2 structures showed two distinct conformations; the open state, where the substrate and cofactor binding sites are accessible to the solvent [found in apo-NATA2 (full and Δ) and NATA2Δ bound to spermine or to di-CoA], and the closed state where substrate binding site and acetyl-containing tail of acCoA are covered by the protein (NATA2Δ bound to acCoA and HEPES) (**Figure 4B**). NATA1Δ structures are all in the closed conformation, although the structure with only acCoA is less firmly closed than the structures with CoA and substrates.

The transition from open to closed involves a pinching movement of the NATA2 regions 39–82 and 196–201 (corresponding to NATA1 residues 32-74 and 188-193), located on opposite sides of the central beta sheet. This movement covers the substrate and catalytic site. Additionally, an outward movement and partial restructuring of residues 161–188 reshape the binding region for the diphosphate moiety of acCoA (NATA1 residues 153-180; **Figure 4B**). Considering the marked thermal stabilisation of NATA1Δ by CoA, it is tempting to speculate that acCoA binding promotes the closed state, at least in NATA1, possibly by acCoA-initiated restructuring of residues 153–180. In support, most of our NATA1 or NATA2 cofactor-bound structures were in a closed state (**Supplementary Table 1**). However, not all cofactor-associated structures of SSATs are closed. For instance, NATA2Δ in complex with the di-CoA adopts the open form, as do some (ac)CoA-bound molecules from mouse, human, and moss structures (PDB 3BJ8, 2B4B chain B, and 7ZHC chain B, respectively). In all these open structures, the CoA diphosphates are bound in the same position as in the closed form, but the adenosine is curled back in the active site (through hydrophobic contacts with helix α1), a position that would clash with the closed form (where the adenosine and 3’ phosphate are extending outside the pocket (**Supplementary Figure 7A**). Conversely, the *Pseudomonas* SSAT crystal structure (PDB 2FE7) shows a closed conformation even in the absence of CoA. We concluded that, although cofactor binding may promote the closed form, the joint presence of the substrate is required. Moreover, the conformation of NATA/SSAT is also strongly influenced by the crystallisation conditions and packing, and/or the energy required for the transition varies between SSATs (we obtained open structures only for NATA2, and not for NATA1).

### Crystal structures suggest catalytic mechanism

In the closed NATA states, the substrate binding site and the acCoA tails (comprising the acetyl moiety) are encapsulated within a tunnel that traverses the protein core (**Figure 4B**). The opening of the substrate side of the tunnel is substantially narrower in the closed NATA1 and NATA2 than in the closed human SSATs.

In the open NATA2Δ structure with spermine (PDB 8XJB), this substrate is lodged in the middle of the catalytic site, on the side of the central β-sheet. In the full-length apo-NATA2 structure (PDB 8RMZ), an ethanediol molecule is found in the same place, mimicking the substrate. This position is partially overlapping with the acCoA binding site (**Figure 5A**). Hence, upon acCoA binding and NATA2 closure, spermine would be pushed toward the back of the substrate binding site, near to the centre of the NATA protein dimer. The position of spermine and ethanediol in the open NATA2 structures is similar to the position of spermine or ethanediol in the open-like mouse and moss SSAT structures (PDB 3BJ8, and 7ZHC_chain B, respectively). In these structures the (ac)CoA tail is curled up to avoid clashes with the centrally bound substrate (mimics). The contracted cofactor form is not compatible with the closed SSAT/NATA conformation. Due to its length, spermine is likely to protrude out of the catalytic pocket in the closed form, as seen for the human SSAT1 bound to BE333 (PDB 2B4B). Conversely, our NATA1Δ structure bound to CoA and ornithine shows that ornithine can be completely captured in the enclosed binding site, similarly to moss SSAT bound to CoA and lysine (PDB 7ZKT).

**Figure 5.**
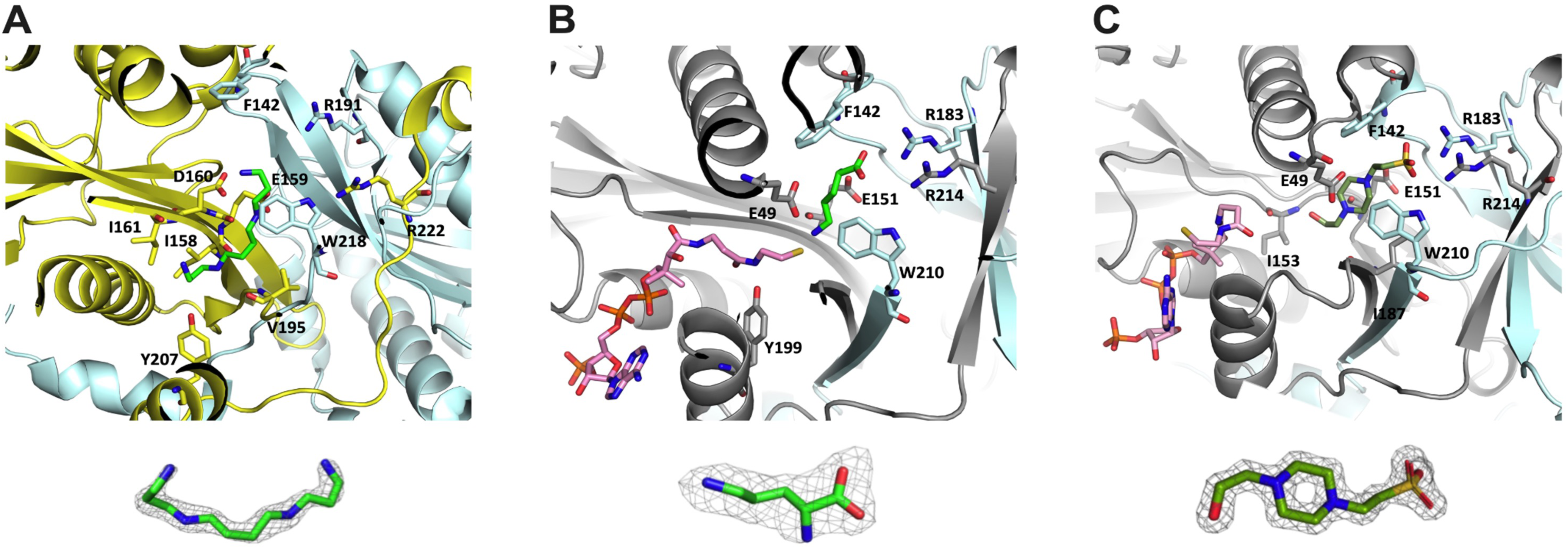
Substrate binding pockets of NATA1Δ and NATA2Δ. **A**) *Top*: Zoom view of the open NATA2Δ structure bound to spermine (green stick model). The two chains of NATA2Δ are shown in yellow and cyan. The side chains involved in binding to spermine are highlighted. *Bottom:* 2Fo-Fc omit map of spermine contoured to 1**σ**. **B**) *Top*: Zoom view of NATA1Δ bound to ornithine (green stick) and CoA (pink stick). The two protein chains are shown in grey and cyan; involved side chains are highlighted. *Bottom*: 2Fo-Fc omit map of ornithine contoured to 1**σ**. **C**) NATA1Δ bound to HEPES (olive green stick) and CoA (pink stick). The di-arginines coordinate the HEPES sulphate group. The catalytic E56 hydrogen-bonds with nitrogen N1 of HEPES, the backbones of E159 and D160 form hydrogen bonds with N4 and O8 of HEPES, and W218 and F150 form hydrophobic interactions with HEPES. Protein chains are shown in grey and cyan, and involved side chains are highlighted. *Bottom*: 2Fo-Fc omit map of HEPES. The omit map is contoured at 1**σ**.

Hence, a reaction cycle would be initiated with substrate binding to the centre of the catalytic pocket (residues I158, E159, D160, Y207, and W218’ in NATA2) where the amine group of spermine directly interacts with OD1 of D160, and W218’ provides a hydrophobic surface for the alkyl groups of spermine (**Figure 5A**). Binding of the acCoA diphosphates to NATA in the presence of the substrate would trigger protein closure, which forces acCoA in an elongated form, pushing the substrate towards the opposite end of the substrate binding pocket. This movement places the tails of the polyamine and acCoA directly under the active site residues (E49, Y199 and E56, Y207 in NATA1 and NATA2, respectively) for transfer of the acetyl moiety (**Figure 5B**).

### The substrate binding pocket of NATA1 and NATA2 appears maladapted to polyamines

The substrate binding pocket of NATA1/2 beyond the active site residues differs markedly from the animal paralogues. In NATA1 and NATA2, the back of the substrate-binding pocket is formed by two arginines, one from each molecule of the dimer, pointing their side chains at the substrate (R183/R214 in NATA1 and R191/R222 in NATA2)(**Figure 5C**). This di-arginine feature is also found in moss SSAT and the bacterial SSATs (e.g. PDB 7ZHC, 2FE7). In the human form, this site is less positively charged. Hence, the substrate binding pocket of plant SSAT/NATAs is more similar to the bacterial one than to the one from animals.

Given that polyamines are positively charged, the di-arginine motif concluding the substrate site is unexpected, as it seems to be adapted to substrates with acidic tails. In support, in the open apo-NATA2 crystal structure (PDB 8RMZ) the di-arginine motif coordinates a sulphate atom from the crystallisation solution (**Supplementary Figure 7B**). In the NATA2Δ structure bound to di-CoA (PDB 8XJH), these arginines coordinate the diphosphate of the second CoA (**Supplementary Figure 7C**). Moreover, NATA1Δ and NATA2Δ structures determined from crystallisation conditions that contained 100 mM HEPES clearly showed one molecule of HEPES bound in their substrate binding site, even though these crystals were exposed to substrates. In both structures, di-arginine side chains coordinate the negatively charged sulphate moiety of HEPES. Additionally, HEPES is tightly bound by hydrophobic and hydrogen bonds (**Figure 5C**), suggesting that HEPES is a potent inhibitor. The inhibitory potential of HEPES was supported by the observation that in the NATA2Δ structure (where HEPES was added before the substrate) the acetyl group of acCoA was intact and extended towards the HEPES molecule. Conversely, in NATA1Δ (where HEPES was added after the substrate), the thiol moiety of acCoA points away from HEPES, and the acetyl group was absent, suggesting it has been used to acetylate substrate prior to exposure to HEPES. Indeed, 5 mM HEPES potently inhibited catalysis of both NATA1Δ and NATA2Δ *in vitro* (**Figure 6A and Supplementary Figure 8 A,B**).

**Figure 6.**
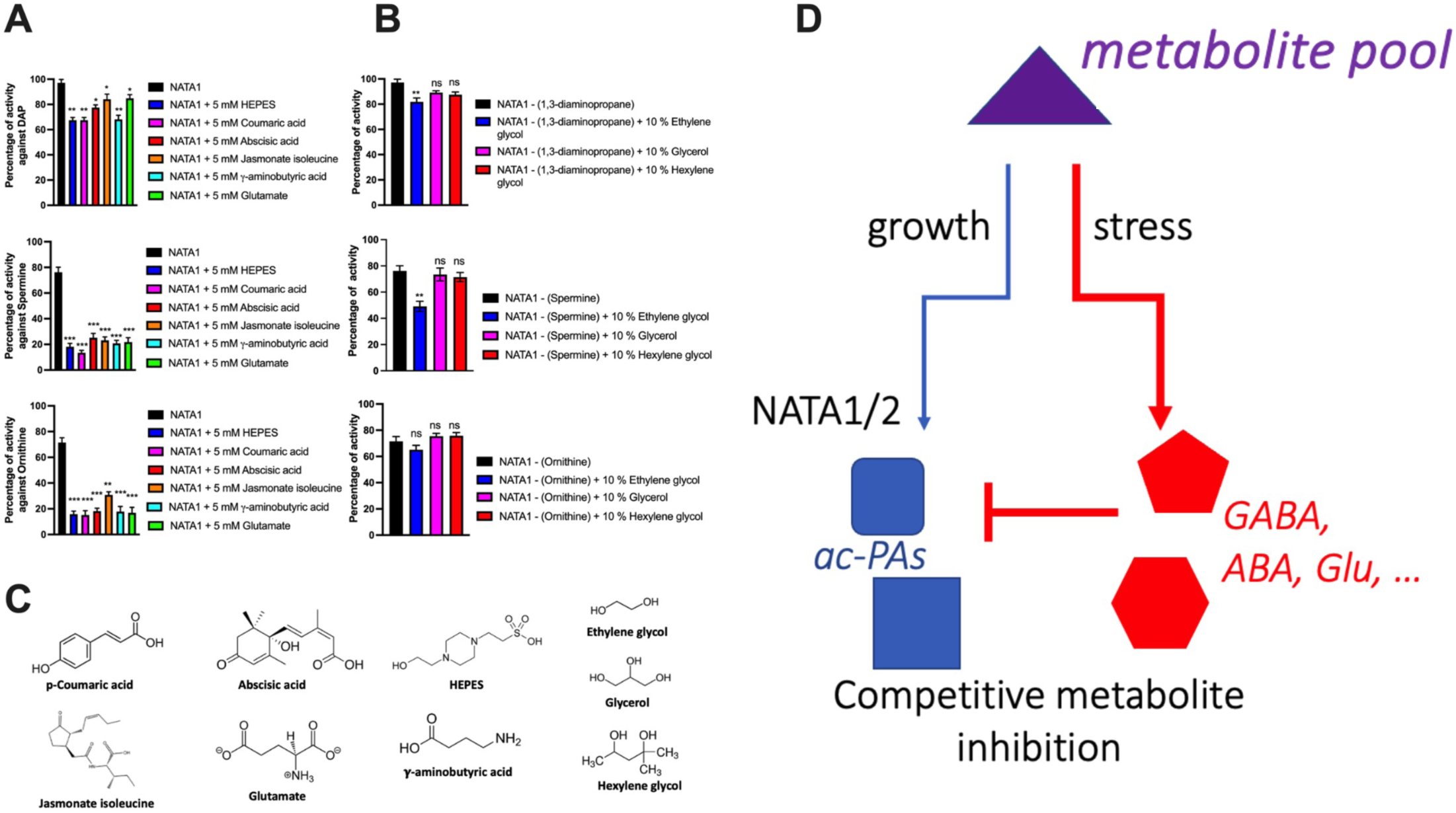
Modulators of NATA1 activity. **A**) Catalytic activity of NATA1Δ against its substrates (from top to bottom: 1,3 diaminopropane, spermine, ornithine), in the presence of HEPES and 5 mM acidic endogenous metabolites in buffer Tris8.5 at 25°C. **B**) Catalytic activity of NATA1Δ against its substrates in the presence of ethylene glycol, glycerol, and hexylene glycol in buffer Tris8.5 at 25°C. **C**) Chemical structure of the HEPES and endogenous metabolites used in the activity assays. **D**) Conceptual drawing for NATA1/2 control by metabolites that reflect the cellular state. *Blue*: Under good growth conditions, NATA1/2 can acetylate the pool of polyamines to maintain homeostasis. *Red*: Acidic metabolites produced under stress conditions can directly inhibit NATA1/2, thus increasing the polyamine concentrations in ways that are favourable for the stressed plant.

Crystallisation trials in the absence of HEPES allowed us to obtain a structure of NATA1Δ in the presence of CoA and ornithine. In this structure, the NATA1 di-arginine motif coordinates the acidic group of ornithine. The N-δ of ornithine was placed between the β-mercaptoethylamine group of CoA and the catalytic residues E49 and Y199, suggesting that this structure positions ornithine and CoA as in the active complex (**Figure 5B**). However, the smaller ornithine appeared more loosely bound in the binding site compared to HEPES, judging by the poorer definition of its 1.35 Å-resolution electron density compared to HEPES. The position of ornithine in closed NATA1Δ corresponds to the position of lysine in moss SSAT (PDB 7ZKT), which also has a di-arginine motif (**Supplementary Figure 7D**).

Given that these structures indicate that substrates need to terminate in a negative charge to match the substrate binding site, it is intriguing that the positively charged diaminopropane, spermine and putrescine are better substrates than ornithine, judged by their Kcat/Km (**Table 2**). All our efforts to co-crystallise spermine and CoA with NATA1 or NATA2 in their closed protein conformation failed, and to date no closed SSAT structure of plant or bacterial SSATs has been obtained, suggesting that polyamines are only residing very briefly in the NATA active site, due to the charge mismatch.

**Table 2.** Catalytic parameters for *E. coli*–produced NATA1Δ and NATA2Δ. Measurements were carried out in Tris8.5 buffer at 25 or 45 °C.

|  | NATA1Δ (25 °C) |  |  | NATA2Δ (25 °C) |  |  | NATA2Δ (45 °C) |  |  |
| --- | --- | --- | --- | --- | --- | --- | --- | --- | --- |
| Ligand | $K_m$<br>(mM) | $K_{cat}$<br>(s<sup>-1</sup>) | $K_{cat}/K_m$<br>(s<sup>-1</sup> mM<sup>-1</sup>) | $K_m$<br>(mM) | $K_{cat}$<br>(s<sup>-1</sup>) | $K_{cat}/K_m$<br>(s<sup>-1</sup> mM<sup>-1</sup>) | $K_m$<br>(mM) | $K_{cat}$<br>(s<sup>-1</sup>) | $K_{cat}/K_m$<br>(s<sup>-1</sup> mM<sup>-1</sup>) |
| Thialysine | 0.54 ± 0.13 | 2.04 ± 0.20 | 3.78 | 0.33 ± 0.08 | 2.20 ± 0.10 | 6.67 | 0.32 ± 0.05 | 3.14 ± 0.19 | 9.82 |
| Diamino propane | 0.44 ± 0.12 | 1.59 ± 0.08 | 3.61 | 0.37 ± 0.07 | 1.46 ± 0.07 | 3.94 | 0.22 ± 0.04 | 1.65 ± 0.08 | 7.5 |
| Spermine | 2.26 ± 0.50 | 1.56 ± 0.10 | 0.69 | 1.62 ± 0.35 | 1.49 ± 0.08 | 0.92 | 0.67 ± 0.13 | 1.31 ± 0.07 | 1.96 |
| Putrescine | 1.90 ± 0.42 | 1.11 ± 0.12 | 0.59 | 2.29 ± 0.48 | 1.04 ± 0.07 | 0.45 | 1.17 ± 0.31 | 1.00 ± 0.08 | 0.85 |
| Ornithine | 4.04 ± 0.82 | 0.64 ± 0.05 | 0.16 | N. D. | N. D. | N.D. | 1.73 ± 0.42 | 0.45 ± 0.13 | 0.26 |
| Lysine | 12.70 ± 3.52 | 0.58 ± 0.09 | 0.05 | 9.48 ± 2.53 | 0.45 ± 0.10 | 0.05 | 5.68 ± 1.25 | 0.63 ± 0.06 | 0.11 |

### Acidic endogenous metabolites modulate NATA catalytic activity

Our results showed that polyamine substrates poorly match the NATA binding pocket in charge and dimensions, whereas HEPES fitted tightly and inhibited NATA activity *in vitro*. We wondered whether endogenous metabolites with similar stereochemistry to HEPES may also inhibit NATA by outcompeting polyamines. Consequently, we tested the similar-sized acidic compounds coumaric acid, abscisic acid, jasmonate-isoleucine, γ-aminobutyric acid (GABA) and glutamate (**Figure 6A**). Indeed, all compounds inhibited the acetylation of ornithine, spermine and diaminopropane by NATA1Δ *in vitro* similarly to HEPES (**Figure 6A**). Acidic compounds also inhibited catalysis by NATA2Δ (**Supplementary Figure 8A**). Unexpectedly, the inhibition of NATA1Δ by HEPES and other acidic compounds was substantial for spermine and ornithine, but only marginal for diaminopropane (**Figure 6A**). Hence, endogenous acidic compounds can potently and selectively modulate substrate acetylation.

### Allosteric pockets may allow more environment-sensing

We observed that even in the closed state, the catalytic pockets of NATA1 and NATA2 displayed large volumes close to the di-Arg motif that were not occupied by the substrate. In the crystal structure of NATA2Δ with HEPES and acCoA (PDB 8XJF), this pocket was filled with a glycerol molecule, presumably from the glycerol used for cryogenic protection. Glycerol was absent from this position in the corresponding NATA1Δ structure, although the crystals were also cryoprotected with glycerol, possibly reflecting slight differences in pocket size. The structures of the NATA orthologues from *P. patens* and *H. sapiens* also had glycerol bound in this position, and additionally an ethanediol molecule was present in *P. patens* (**Supplementary Figure 8C, Supplementary Table 1**). These structures suggested that the large cavities adjacent to the NATA substrate binding sites may influence catalysis.

, we tested the influence of ethylene glycol, glycerol and hexylene glycol on the activity of NATA1Δ on acetylating diaminopropane, spermine and ornithine. Only ethylene glycol significantly reduced the catalytic activity of NATA1Δ, and only for spermine and ornithine (**Figure 6B**). Glycerol and hexylene showed no effects under these conditions, which was reassuring considering that 10–20% glycerol is frequently used in SSAT reactions and protein buffers [e.g. see (Kuninger et al., 2007; Lou et al., 2016; Toleman et al., 2004)]. The absence of an effect for glycerol also confirmed that the reduced catalysis observed between buffers Tris8.5 and Tris7.5 was due to their different pH (pH 7.5 and pH 8.5) rather than the presence of glycerol in Tris7.5 (**Figure 6C**). However, our results supported that ethylene and possibly other small compounds may selectively influence catalysis by lodging in a crevice next to the active site.

## DISCUSSION

In this study, we integrated *in planta* analyses with structural and biochemical approaches to investigate the biological role and regulation of NATA1 and NATA2. By determining eight structures of Arabidopsis NATA1 and NATA2, we gained unparalleled insights into the conformational transitions associated with substrate catalysis. We showed that NATA2 is a heat-stable isoform of NATA1, explaining why NATA2 is expressed under heat stress. We confirmed that the plant-specific 35-residue insertion is partially structured in the Arabidopsis SSATs, adding an extra strand to the central β-sheet. Although the role of this insertion is still uncertain, our observation of a crystal contact-promoted rearrangement in this region suggests it may function as a ligand binding site.

Our *in planta* analysis of NATA2 echoes previous findings on NATA1, showing that the presence of each enzyme reduces Arabidopsis’ ability to simultaneously maintain growth and stress resistance (Lou et al., 2016). This phenotype aligns with observations that polyamines generally enhance stress resistance in plants (Gill and Tuteja, 2010; Takahashi and Kakehi, 2010; Jang et al., 2012), implying that enzymes reducing polyamines are disadvantageous under stress conditions.

Considering the unfavourable contribution of NATA activity to plant resistance, the NATA gene duplication and the expression of the heat-stable NATA2 paralogue under heat stress is intriguing. However, our analysis also shows that NATA activity is required for seed germination, suggesting that NATA1 and NATA2 perform essential functions that cannot be shut down completely. Notably, NATA1 expression appears to be relatively stable under various abiotic stress conditions, raising the question of how the acetyltransferase function of NATA1 and NATA2 is regulated.

Our research now suggests that Arabidopsis has evolved regulatory mechanisms that operate at the metabolite level (**Figure 6D**). The structural analysis reveals that the substrate binding sites of both NATA1 and NATA2 possess stereochemical characteristics (size and charge), including a di-arginine motif, that seem maladapted for binding to positively charged polyamines. Based on the serendipitous observation that HEPES binds firmly to this pocket and strongly inhibits substrate acetylation, we identified several endogenous plant metabolites that selectively reduced the acetylation of substrates. Of those compounds, coumaric acid, abscisic acid, jasmonate-isoleucine, GABA and glutamate resemble HEPES in terms of size, centrally located hydrophobic region, and negatively charged tail. Hence, we propose that these metabolites bind NATAs in the same way as HEPES, thereby blocking substrate access. These compounds selectively blocked acetylation of spermine and of the negatively charged ornithine (which poorly filled the substrate cavity in our crystal structures). However, the catalysis of the small diaminopropane was only slightly affected.

Interestingly, coumaric acid, abscisic acid, jasmonate-isoleucine, GABA, and glutamate are produced in increasing quantities under stress conditions, where they confer beneficial effects on the plant. Notably, GABA has a range of protective functions that increase plant stress tolerance by improving photosynthesis, inhibiting reactive oxygen species (ROS) generation, activating antioxidant enzymes, and regulating stomatal opening in drought stress (Priya et al., 2019; Sita and Kumar, 2020; Hasan et al., 2021; Guo et al., 2023). GABA plays a role in plant development (including reproduction, germination, root growth, and fruit ripening), and accumulates in response to both biotic stress (such as viruses, bacteria, fungi, or insects) and abiotic stress (including temperature, drought, salt, and ROS)(Li et al., 2021). GABA is produced from glutamate *via* the GABA-shunt pathway, particularly under abiotic stress conditions. Glutamate itself also participates in pathogen resistance, response and adaptation to abiotic stress (such as salt, cold, heat, and drought) (Qiu et al., 2020). Similarly, abscisic acid and jasmonoyl-isoleucine are key regulators of plant growth and defence in response to various biotic and abiotic stress signals. Abscisic acid–deficient mutants from various plant species exhibit reduced seed dormancy and wilted phenotypes (Kermode, 2005; Fujii and Zhu, 2009; Nambara, 2017). Jasmonoyl-isoleucine levels increase in response to wounding or other forms of tissue damage(Suza and Staswick, 2008; Thurow et al., 2020), and *NATA1* is one of the most jasmonate-induced genes in *A. thaliana* (Adio et al., 2011). Coumaric acid is one of several compounds in the hydroxycinnamic acid amide pathway, which play central roles in plant-pathogen interactions (Macoy et al., 2015; Liu et al., 2022).

Given the protective roles of polyamines in Arabidopsis and other plants, our findings suggest that NATA1 and NATA2 have evolved a substrate binding pocket that enables acidic compounds produced under stress to inhibit polyamine acetylation. This mechanism would help to maintain the pool of stress-mitigating polyamines. Here it is particularly interesting that these acidic compounds do not inhibit the acetylation of diaminopropane, because it is its acetylated form which is beneficial for plants under stress conditions (Jammes et al., 2014). Therefore, for Arabidopsis at least, our results propose that the seemingly suboptimal architecture of the catalytic site has evolved to allow a group of stress-produced metabolites to collectively inhibit the acetylation of polyamines needed for stress resistance, while permitting the acetylation of diaminopropane, a modification that is advantageous under stress conditions.

We also discovered that ethylene acts as a selective inhibitory agent, though its potency was low. Our analysis of available crystal structures suggests that this small compound exerts an allosteric inhibitory effect on the catalysis by occupying the space adjacent to the substrate binding site. Ethylene, a plant hormone, is implicated in numerous physiological processes, including plant growth, development, and senescence (Abeles et al., 1992). It also plays a crucial role in plant responses or adaptation to biotic and abiotic stress conditions (Müller and Munné-Bosch, 2015). The production of ethylene often enhances plant tolerance of sub-optimal environmental conditions. In our experiments, ethylene primarily reduced the acetylation of spermine, but had only a marginal effect on diaminopropane and no effect on ornithine. This observation suggests an additional pathway for allosteric effectors of NATA activity in Arabidopsis. Our results suggest that some of the reported discrepancies in the substrate profile and activity of SSATss could be attributed to their biologically relevant sensitivity to a wide array of compounds and conditions. However, other mechanisms, such as post-translational modifications, may also be involved.

In conclusion, our research identifies the molecular basis for the temperature-specific roles of NATA1 and NATA2 paralogues in Arabidopsis, elucidates the substrate turnover mechanism of plant SSATs/NATAs, and proposes that NATA function is regulated under stress conditions through a competitive metabolite-ensemble inhibition mechanism. This mechanism could be more widely employed in plants as a tool for rapid, environment-responsive, non-genetic enzyme regulation.

## Methods

### Plants and growth conditions

Seeds of wild-type Columbia-0 Arabidopsis, *nata2* (Salk_092319, (Alonso et al., 2003), and *nata1* (GK-256F07, (Rosso et al., 2003)) were obtained from the Arabidopsis Biological Resource Center (www.arabidopsis.org). Plants were grown in Cornell mix (by weight 56% peat moss, 35% vermiculite, 4% lime, 4% Osmocoat slow-release fertiliser [Scotts, Marysville, OH], and 1% Unimix [Scotts]) in 20x40-cm nursery flats in Conviron growth chambers with a photosynthetic photon flux density of 200 mmol m^-1^ s^-1^ and a 16:8 h day :night photoperiod, at 23°C with a 50% relative humidity. For experiments carried out at elevated temperature, a chamber set to identical conditions except for a 28°C temperature was used.

### Methyl jasmonate induction

Methyl jasmonate induction is performed as described previously (Lou et al., 2016), with slight modification. In short, the leaves of 4-week-old Arabidopsis plants were sprayed with an aqueous solution containing 0.01% (v/v) Tween 20, and 0.01 mM methyl jasmonate. Control plants were treated with water containing 0.01% Tween 20 and 0.03% acetone, the solvent used for methyl jasmonate. Plants were covered with plastic domes and tissue was harvested 4 days after elicitation and immediately frozen in liquid nitrogen.

### Quantification of polyamines

Polyamines were extracted and derivatized with 6-aminoquinolyl-N-hydroxysuccinimidyl carbamate (AQC) using the AccQ-Fluor reagent kit (Waters) prior to HPLC detection, as described in (Lou et al., 2016). For better sensitivity and further confirmation on expected compounds, a 4-(N,N-dimethylaminosulfonyl)-7-fluoro-2,1,3-benzoxadiazole (DBD-F) derivatization method modified from Lou et al. 2016 was used prior to HPLC-MS detection. In short, plant tissue was pulverised in liquid nitrogen and extracted with 25 mM HCl (5 ml mg^-1^ of tissue) containing 25 μM 1,6-hexanediamine as an internal standard. After centrifuging at 15,000 x g at 10°C for 20 min, the supernatants (12 μL each) were mixed with 24 μL of 0.2 M sodium tetraborate. Equal volumes of DBD-F solution, dissolved in acetonitrile, were added to each sample prior to 30 min incubation at 60°C. The reaction mixtures were filtered and analysed on a Waters^®^ ACQUITY UPLC^®^ BEH C18 column (Waters) using a Thermo Q-Exactive Orbitrap coupled to Dionex ultra-high-pressure chromatography systems for reverse-phase separation with column temperature set at 40 °C. The starting conditions were 80% solvent A (0.1% formic acid in water) and 20% solvent B (acetonitrile) with flow rate at 0.5 mL/min. The elution gradient was as follows: ramp to 90% solvent B at 5.5 min with curve set to 5 and hold at 90% solvent B until 7 min. The detection conditions were as follows: Heated electrospray ionisation mode; 4000 V spray voltage; 320°C capillary temperature; 70 units sheath gas flow rate; mass range of m/z 80 - 1200 Da. The analytical software Therm Xcalibur v3.0 was used for the system control and data processing.

### Imbibed seed experiment

Fifty mg of wild type, *nata1* and *nata2* mutant seeds were imbibed in sterile distilled water for 0 hr, 24 hr, and 48 hr. Seeds were homogenised and extracted as described above.

### Metabolite profiling

For non-targeted metabolomic assays, plants were grown and treated with methyl jasmonate, as described above. Samples were separated on an Acclaim (Thermo Scientific) column using a Dionex Ultimate 3000 UHPLC system, and metabolites were detected using a quadrupole time-time-of flight mass spectrometer (MicrOTOF-Q II; Bruker Daltronics), as described previously (Tzin et al., 2015). Raw mass spectrometry data files from positive and negative ionisation modes were processed in R using the XCMS (http://metlin.scripps.edu/download/) and CAMERA (http://www.bioconductor.org/packages/release/bioc/html/CAMERA.html). Processed data sets with a total of 5932 mass features were transferred to Microsoft Excel prior to PLS-DA analysis using Metaboanalyst 3.0 (http://www.metaboanalyst.ca/faces/home.xhtml; (Xia and Wishart, 2016), with the following program parameters: upload data = peak intensity table, missing value estimation = skip, data filtering = mean intensity value, data transformation = logarithmic, and data scaling = auto scaling.

### Heat stress and hypocotyl elongation assay

To test the sensitivity of hypocotyl elongation to heat stress, an experiment similar to that described by (Shen et al., 2016), was performed. In short, seeds were germinated on half-strength Murashige and Skoog medium (½ MS salts, 1.5% sucrose, pH 5.7) with 1.2% agar and left to grow in a vertical position in a dark chamber (23°C). After three days, the plates were sealed and immersed in a 37°C water bath for three hours before transferring back to the dark chamber for another three days of growth in a vertical position. Photographs were taken with a Nikon D80 digital camera, and the length of the hypocotyls was measured with ImageJ (www.imagej.net/). For polyamine analysis after heat stress, half-strength MS plates with 7-day-old seedlings were sealed and immersed in a 37°C water bath for three hours. Twelve to 16 seedlings from the same plate were combined and ground to a fine powder in liquid nitrogen for polyamine extraction. For quantitative PCR analysis, a similar experiment was performed in a staggered manner, so that seedlings exposed to heat stress for 0 hr, 1 hr, and 2 hr were harvested at the same time.

### Quantitative PCR

Relative transcript abundances of *NATA2* (At2g39020), *HSP70* (At5g02500), *NATA1* (At2G39030), *PR1* (At2g14610), *PR2* (At3g57260), and *PR5* (At1G75040) were analysed and compared by qRT-PCR using actin (At3g18780) as an endogenous control as described in Lou et al. 2016. Gene-specific primers were designed using Primer-Blast (www.ncbi.nlm.nih.gov/tools/primer-blast/) and listed in **Supplementary Table S4**. Rosette leaf samples were ground in liquid nitrogen and total RNA was extracted using the SV total RNA isolation system (Promega, www.promega.com). After quantification with a Nanodrop system (www.nanodrop.com), 1 μg of total RNA was reverse transcribed using SMART^®^ MMLV Reverse Transcriptase (Clontech, www.clontech.com) using oligo(dT)15 primers. Following cDNA synthesis, the samples were diluted in nuclease free water and used for qRT-PCR reactions with the SYBR^®^ Green PCR master mix (Applied Biosystems, www.appliedbiosystems.com) using an Applied Biosystems 7900HT Instrument. Each reaction was carried out with the following conditions: 95°C for 10 min, followed by 40 cycles of 95°C for 15 s, 60°C for 15 s and 72°C for 15 s, and final extension at 72°C for 2 min. The CT values were quantified and analysed according to the standard curve method.

### Bacterial growth assays

An infection experiment with *Pseudomonas syringae* strain DC3000 was performed as described by (Lou et al., 2016). In short, bacteria were cultured at 30°C in Luria-Bertani (LB) medium supplemented with 50 μg ml^-1^ rifampicin and 50 μg ml^-1^ kanamycin. Overnight cultures were centrifuged at 3,000 x g for 10 min, resuspended, and diluted in water to approximately 10^5^ colony forming units (CFU) ml^-1^ before infiltration into three-week-old Arabidopsis rosette leaves with a needleless syringe. Eight-mm diameter leaf discs were collected 2 days after infiltration and submerged in 1 ml sterile water with 0.01% Tween 20 for 4 h to allow equilibration of the bacteria between the apoplastic space of the leaf disks and the surrounding water, as described previously (Adio et al., 2011). Serial dilutions of the suspensions were spotted on LB agar plates supplemented with 50 μg ml^-1^ rifampicin, and bacterial colonies were counted after 2 days of incubation at 30°C.

### Screening for double mutant lines

Heterozygous *nata2* mutations in a homozygous *nata1* mutant background were made by TALEN-induced mutagenesis (Christian et al., 2013) (**Supplementary Figure 2B**). Progeny from self-pollinated *nata1/nata1 NATA2/nata2* plants were genotyped. DNA was extracted using the CTAB (cetyltrimethylammonium bromide) method (Doyle, J.J. and Doyle, J.L., 1987), amplified by PCR, and digested with restriction enzyme NspI, which cuts in the wild-type *NATA2* gene but not in the *nata2* deletion mutant. Digested bands were visualised on 1.5% agarose gels. Absence of the hygromycin resistance gene was screened to segregate away the TALEN construct.

### Cloning, expression, and purification of NATA 1/2

N-terminally truncated NATA1 (19-228, NATA1Δ) and NATA2 (30-236, NATA2Δ) as well as the full-length NATA2 from Arabidopsis were cloned to the pGEX6P-1 expression vector (GE Healthcare). The plasmids were then transformed into Rosetta 2 (DE3) cells for expression. Cells producing the two truncated forms were grown in LB medium containing ampicillin (100 μg/ml) and chloramphenicol (35 μg/ml) at 37°C until an OD600 of 0.6. Protein expression was induced by adding 150 μM isopropyl β-D-1-thiogalactopyranoside (IPTG), and the cultures were incubated at 16 °C overnight. For the full-length production, cells were grown in Terrific Broth medium supplemented with 200 mg/L carbenicillin and 30 mg/L chloramphenicol. Protein expression was induced at OD600 of 0.8 by adding 100 µM IPTG, and the cells were cultured at 18°C, overnight. Further affinity purification steps were followed as previously reported by (Shahul Hameed et al., 2022). After GST cleavage, the eluted proteins were purified on a HiLoad16/60 Superdex 200 prep-grade gel filtration column (GE Healthcare) using a buffer containing 20 mM Tris (pH 7.5), 150 mM NaCl, and 3 mM DTT. Protein purity was evaluated using SDS-PAGE. The purified NATA1Δ and NATA2Δ as well as the full-length NATA2 were concentrated to 10 mg/ml, 4 mg/ml and to 5.2 mg/ml respectively, and stored at -80° C.

### Crystallisation and structure determination

NATA1Δ and NATA2Δ were preincubated with 2 mM acCoA (Roche) and 5 mM of ligands (purchased from Sigma) such as ornithine, spermine, and γ-amino butyric acid. The protein complexes were then subjected to the sitting drop vapour diffusion method for crystallisation screening using commercially available sparse matrix screens. NATA1Δ crystals were obtained in the reservoir solutions containing 0.2 M Magnesium chloride hexahydrate, 0.1 M HEPES pH 7.0, 20% w/v PEG 6000, and 0.2 M Ammonium sulphate, 0.1 M Tris pH 8.5, 12% w/v PEG 8000. NATA2Δ crystals were obtained in the reservoir solution containing 0.2 M Sodium citrate tribasic dihydrate, 0.1 M Tris pH 8.5, 30% v/v PEG 400, and, 0.2 M Potassium thiocyanate 0.1 M Bis-Tris propane pH 7.5, 20% w/v PEG 3350, and, 0.2 M Lithium chloride, 0.1 M HEPES pH 7.0, 20% w/v PEG 6000. The crystals appeared after 7 days at 23 °C. Apo NATA2 (full-length) crystals were obtained in the reservoir solution containing 0.2 M Ammonium sulphate, 0.1 M Na-citrate pH 5.6, 15% v/v PEG 4000. As crystals were only identified after more than four weeks, a partial proteolysis of the flexible N-terminal residues 1-19 cannot be excluded. For data collection, 25% glycerol was used as a cryo-protectant and the crystals were flash-cooled in liquid nitrogen. Data were collected at 100 K at the beamlines PROXIMA 1 and PROXIMA 2A at the SOLEIL Synchrotron (France), using EIGER X 16M and EIGER X 9M detectors, respectively (Proposal numbers 20191181, 20201179, 20210195, 20210932, 20220373). The data were processed using either the “XDS made easier” pipeline (Legrand et al., 2019) or autoPROC (www.globalphasing.com) including STARANISO (Vonrhein et al., 2011). Initial phases were determined using human thialysine acetyltransferase (SSAT2) structure (PDB: 2BEI) as a search model for molecular replacement using Balbes or Phaser. The structures were manually inspected and corrected using Coot and refined using either Phenix Refine or BUSTER-2.10.4 (Blanc et al., 2004) using TLS group and NCS restraints (**Supplementary Table 2**). The figures were drawn with PyMOL.

### Thermal shift assay

The thermal stability of NATA1Δ and NATA2Δ in the presence and absence of different ligands and cofactors was determined using thermal unfolding profiles. All assays were carried out at a final protein concentration of 10 μM, in buffer containing 50 mM Tris–HCl pH 8.5, 150 mM NaCl, and with Sypro Orange dye (Invitrogen) at a 2× final concentration. Thermal denaturation temperatures were obtained in a CFX96 touch real-time thermal cycler (Bio-Rad) with an optical reaction module by ramping the temperature between 20 and 95 °C at 1 °C per minute, and fluorescence acquisition through the channel. Data were analysed with CFX Manager Software V3.0 (Bio-Rad) and plotted using PRISM version 9.0 (www.graphpad.com).

### Acetyltransferase activity using DTNB assay

The catalytic activity of NATA1Δ and NATA2Δ was measured spectrophotometrically on a Pherastar plate reader (BMG Labtech) at 25 °C using dithiobis-(2-nitrobenzoic acid) (DTNB) assay. The assay was carried out using various ligands following the previously reported work by Belicek et al., 2023. The substrate screening was made in 50 mM Tris–HCl pH 8.5, 150 mM NaCl, using 0.5 acCoA, 5 mM substrate, and 0.5 mM DTNB. The data were plotted using PRISM version 9.0 (www.graphpad.com).

### Isothermal Titration Calorimetry

NATA1Δ and NATA2Δ were dialysed and degassed in the isothermal titration calorimetry (ITC) buffer (20 mM Tris pH 7.5, 150 mM NaCl, 3 mM DTT). AcCoA (Roche) and CoA (Sigma) were dissolved in the identical buffer. For the titrations, 20 μM of NATA1Δ and NATA2Δ were placed in the cell, and 350 μM of acCoA and CoA were loaded in the syringe. Titrations were performed at 25 °C with an initial injection of 0.4 μl, followed by 18 injections of 2 μl. ITC experiments were performed on ITC200 (Malvern Panalytical, UK). Data analysis was performed and figures prepared using MicroCal Origin 7 software.

### Declaration of generative AI and AI-assisted technologies in the writing process

During the preparation of this work the authors used Copilot in order to improve the language in some passages. After using this tool/service, the authors reviewed and edited the content as needed and take full responsibility for the content of the publication.

## Supporting information

datset

## Acknowledgments

This research was funded by US National Science Foundation awards 1121788 and 1022017 to GJ and by King Abdullah University of Science and Technology (KAUST) through the baseline fund to STA. This work benefited from the I2BC crystallisation platforms supported by FRISBI ANR-10-INSB-05-01. We acknowledge SOLEIL for provision of synchrotron radiation facilities and would like to thank A. Thompson, M. Pierre, P. Legrand, S. Sirigu, M. Savko and B. Shepard for assistance in using the beamlines PROXIMA 1 and PROXIMA 2A (Proposal IDs: 20191767, 20200610, 20201179, 20210932, and 20230217). For computer time, this research used the resources of the KAUST Supercomputing Laboratory, and experimental research was supported by the Bioscience Core Lab and the Imaging and Characterization Core Lab at King Abdullah University of Science & Technology (KAUST) in Thuwal, Saudi Arabia. We thank Navid Mohaved for assistance with mass spectrometry assays and Yiping Qi for providing the *nata2* TALEN knockout line described by Christian et al 2013. E. Aleksenko thanks V. Gaudin and J. Leung for supervision, Y. Sulio for his help, and the University of Paris-Saclay for her PhD fellowship.

## Conflict of Interest Statement

The authors have no conflicts of interest related to the research in this manuscript.

## Author Contributions

YL, JY, and GJ conceived the plant experiments; YL and JY conducted plant experiments; UFSH, and STA conceived the biochemical and biophysical analyses, and UFSH conducted these experiments. EA produced the full-length NATA2 protein. UFSH, STA, PB, and SM determined and analysed the protein structural data. STA, YL and GJ wrote the manuscript. All authors read, commented on, and approved the manuscript.

## Figure Legends

**Supplementary Figure S1. Comparison of SSAT homologues. A**) Multiple sequence alignment of the SSATs from different species. The plant SSAT–specific loop sequences are boxed in red. The additional basic residues uniquely present in the P-loops of plant SSATs are boxed in blue. The two arginines from the di-arginine motif in the catalytic pocket are marked by an asterisk (*), and the three residue positions important for the heat stability of NATA2 versus NATA2 are marked by a cross (✣). Secondary structure derived from NATA1 is shown above the sequences. **B**) Superimposition of the PDB structures of SSAT dimers. NATA1Δ - grey; NATA2Δ - yellow; *Pp*SSAT (PDB 7ZHC) - blue; *Hs*SSAT1 (PDB 2B3U) - salmon; *Hs*SSAT2 (PDB 2BEI) - green; *Mm*SSAT1 (PDB 3BJ8) - purple. The additional loop present in plant SSATs is shown in red.

**Supplementary Figure S2. Design and effects of nata2 mutant plants. A**) Accumulation of the polyamines ornithine (*left*), putrescine (*middle*) and *N*-acetylputrescine (*right*) in rosette leaves of wild-type *A. thaliana* and *nata2* mutant plants, as measured by HPLC. Mean ± SE of N=9. **B**) NATA2 sequence changes due to the TALENS. (*Top*) Schematic overview of the strategy to disrupt *NATA2* in the *nata1* mutant background. RVD, Repeat Variable Diresidue. (*Bottom*) Sanger sequencing result of the selected line used for generating a *nata1 nata2* double mutant. **C**) Spatial and temporal expression of *NATA2,* At2g39020. Data are from ePlant (https://bar.utoronto.ca/eplant/; Waese et al. 2017). **D**) *A. thaliana nata2* seedlings show no obvious variation in the growth phenotype compared to Col-0 wild-type seedlings after two and five days growth on half-strength Murashige and Skoog agar plates.

Supplementary Figure S3. *NATA2* gene expression in response to abiotic stress. http://jsp.weigelworld.org/expviz/; (Schmid et al., 2005).

**Supplementary Figure S4. NATA1Δ catalytic activity in the presence of additives (ac)CoA binding. A)** Comparison of the catalytic activity of NATA1Δ against its substrates in the presence of ethylene glycol, glycerol, and hexylene glycol in buffer Tris8.5 and Tris7.5 at 25°C. **B**) ITC investigating binding of (ac)CoA to NATA1Δor NATA2Δ. *Top panels*: raw data showing heats for each injection. *Bottom panels*: Integrated heats from top panel. **C**) Table of binding affinities and the thermodynamic parameters calculated from (B).

**Supplementary Figure S5. Structural details of NATA1Δ/NATA2Δ.** A) Zoom view into the plants-specific insert and its interactions with the conserved NATA core. Shown are superimposed structures of NATA1Δ bound to HEPES and CoA (grey), NATA2Δ bound to HEPES and acCoA (yellow) and NATA2Δ bound with di-CoA (magenta), produced in-cristallo. Key residues involved in the interaction between the loop and core residues are shown as sticks. **B**) Structural rearrangement of the β-strand from the plant-specific insert caused by a symmetry-related molecule (orange) in the NATA2Δ bound to di-CoA crystal structure. The inlay zooms into the displaced β-strand of the di-CoA–bound NATA2Δ structure (marked by an arrow). Superimposed is the structure of NATA2Δ bound to HEPES-CoA (yellow) with the unaffected β-strand conformation. **C**) Position of the additional arginine present in the P-loop of higher plants, including Arabidopsis. Arginines are shown as stick models (NATA1Δ: grey carbons; NATA2Δ: yellow carbons). The acCoA is shown as a stick model with green carbons.

**Supplementary Figure S6. Molecular basis of NATA2 heat stability and its validation by mutations. A**) Superimposition of NATA1Δ bound to HEPES and CoA (grey) and NATA2Δ bound to HEPES and acCoA (yellow). Zoom figure shows the central core residues identified as modulating the stability of the proteins. **B**) Table showing the heat stability values of NATA1Δ and NATA2Δ and its mutants using the FoldX server. All values are in kcal/mol. **C**) DSF traces showing the heat stability of NATA1Δ and NATA2Δ and mutants.

**Supplementary Figure S7. Details of NATA1Δ, NATA2 and NATA2Δ and their ligand interactions. A**) Zoom view of NATA2Δ (magenta) bound to di-CoA (blue carbons). di-CoA and NATA2Δ residues involved in binding are shown as sticks. **B**) Zoom view of the structure of apo-NATA2 (yellow) crystallised from a solution containing full-length protein. Sulphate atoms shown as sphere models and the NATA2 residues are shown as sticks. The ethanediol (EDO) bound in the catalytic pocket is shown as a stick model with carbons in yellow. **C**) 2Fo-Fc omit map of di-CoA contoured at 1**σ**. **D**) Superimposition of NATA1Δ bound to ornithine and CoA (grey), and of *Pp*SSAT bound to lysine and acCoA (pdb: 7zkt; blue). Stick models show ornithine (green carbons), lysine (yellow), acCoA (pink) and the di-arginines involved in binding.

**Supplementary Figure S8. Additional experimental and structural data. A**) Catalytic activity of NATA2Δ on spermine in the presence of acidic compounds. **B**) DSF data indicating heat stability of NATA1Δ and NATA2Δ in the presence of acidic endogenous metabolites. **C**) Glycerol and ethanediol bound in the catalytic pocket are shown as sticks, colour-coded according to the crystal structure in which they were identified: NATA2Δ bound to HEPES and CoA (yellow), full-length apo-NATA2 (orange), and *Pp*SSAT (blue; pdb 7zkt).

**Supplementary Table 1. Summary of NATA/SSAT structures.**

**Supplementary Table 2. Crystallographic data.**

**Supplementary Table 3A/B. Thermal melting point (*Tm*) of NATA1Δ and NATA2Δ established by DSF.** Analysis was carried out in buffer Tris8.5.

**Supplementary Table 4A/B. Primers used A)** Primers used for cloning and genotyping. **B**) Primers used for quantitative RT-PCR.

**Supplemental Data Set S1. HPLC-TOF-MS analysis.** Analysis of wild-type Columbia-0, nata1, and nata2, with and without methyl jasmonate treatment (raw metabolomic data shown used PLS-DA analysis in Figure 1E.)

## Notes

### Competing Interest Statement

The authors have declared no competing interest.

## References

Abeles, F. B., Morgan, P. W., and Saltveit, J. Ethylene in Plant Biology. 2nd ed.

Adio, A. M., Casteel, C. L., De Vos, M., Kim, J. H., Joshi, V., Li, B., Juéry, C., Daron, J., Kliebenstein, D. J., and Jander, G. (2011). Biosynthesis and Defensive Function of *N* Δ-Acetylornithine, a Jasmonate-Induced *Arabidopsis* Metabolite. The Plant Cell 23:3303–3318.

Alonso, J. M., Stepanova, A. N., Leisse, T. J., Kim, C. J., Chen, H., Shinn, P., Stevenson, D. K., Zimmerman, J., Barajas, P., Cheuk, R., et al. (2003). Genome-Wide Insertional Mutagenesis of *Arabidopsis thaliana*. Science 301:653–657.

Bagni, N., and Tassoni, A. (2001). Biosynthesis, oxidation and conjugation of aliphatic polyamines in higher plants. Amino Acids 20:301–317.

Bělíček, J., Ľuptáková, E., Kopečný, D., Frömmel, J., Vigouroux, A., Ćavar Zeljković, S., Jagic, F., Briozzo, P., Kopečný, D. J., Tarkowski, P., et al. (2023). Biochemical and structural basis of polyamine, lysine and ornithine acetylation catalyzed by spermine/spermidine *N* -acetyl transferase in moss and maize. The Plant Journal 114:482–498.

Bewley, M. C., Graziano, V., Jiang, J., Matz, E., Studier, F. W., Pegg, A. E., Coleman, C. S., and Flanagan, J. M. (2006). Structures of wild-type and mutant human spermidine/spermine *N* ^1^ -acetyltransferase, a potential therapeutic drug target. Proc. Natl. Acad. Sci. U.S.A. 103:2063–2068.

Blanc, E., Roversi, P., Vonrhein, C., Flensburg, C., Lea, S. M., and Bricogne, G. (2004). Refinement of severely incomplete structures with maximum likelihood in *BUSTER– TNT*. Acta Crystallogr D Biol Crystallogr 60:2210–2221.

Cermak, T., Doyle, E. L., Christian, M., Wang, L., Zhang, Y., Schmidt, C., Baller, J. A., Somia, N. V., Bogdanove, A. J., and Voytas, D. F. (2011). Efficient design and assembly of custom TALEN and other TAL effector-based constructs for DNA targeting. Nucleic Acids Research 39:e82–e82.

Christian, M., Cermak, T., Doyle, E. L., Schmidt, C., Zhang, F., Hummel, A., Bogdanove, A. J., and Voytas, D. F. (2010). Targeting DNA Double-Strand Breaks with TAL Effector Nucleases. Genetics 186:757–761.

Christian, M., Qi, Y., Zhang, Y., and Voytas, D. F. (2013). Targeted Mutagenesis of *Arabidopsis thaliana* Using Engineered TAL Effector Nucleases. G3 Genes|Genomes|Genetics 3:1697–1705.

Coleman, C. S., Hu, G., and Pegg, A. E. (2004). Putrescine biosynthesis in mammalian tissues. Biochemical Journal 379:849–855.

Della Ragione, F., and Pegg, A. E. (1983). Effect of analogues of 5′-methylthioadenosine on cellular metabolism. Inactivation of *S* -adenosylhomocysteine hydrolase by 5′-isobutylthioadenosine. Biochemical Journal 210:429–435.

Doyle, J.J. and Doyle, J.L. (1987). A Rapid DNA Isolation Procedure for Small Quantities of Fresh Leaf Tissue. Phytochemical Bulletin. Phytochemical Bulletin 19:11–15.

Evans, P. T., and Malmberg, R. L. (1989). Do Polyamines Have Roles in Plant Development? Annu. Rev. Plant. Physiol. Plant. Mol. Biol. 40:235–269.

Fliniaux, O. (2004). Altered nitrogen metabolism associated with de-differentiated suspension cultures derived from root cultures of Datura stramonium studied by heteronuclear multiple bond coherence (HMBC) NMR spectroscopy. Journal of Experimental Botany 55:1053–1060.

Fujii, H., and Zhu, J.-K. (2009). Arabidopsis mutant deficient in 3 abscisic acid-activated protein kinases reveals critical roles in growth, reproduction, and stress. Proc. Natl. Acad. Sci. U.S.A. 106:8380–8385.

Galston, A. W., and Sawhney, R. K. (1990). Polyamines in plant physiology. Plant Physiol. 94:406–410.

Galston, A. W., Kaur-Sawhney, R., Altabella, T., and Tiburcio, A. F. (1997). Plant Polyamines in Reproductive Activity and Response to Abiotic Stress*. Botanica Acta 110:197–207.

Gill, S. S., and Tuteja, N. (2010). Polyamines and abiotic stress tolerance in plants. Plant Signaling & Behavior 5:26–33.

Guo, Z., Gong, J., Luo, S., Zuo, Y., and Shen, Y. (2023). Role of Gamma-Aminobutyric Acid in Plant Defense Response. Metabolites 13:741.

Han, B. W., Bingman, C. A., Wesenberg, G. E., and Phillips, G. N. (2006). Crystal structure of *Homo sapiens* thialysine *N* ^ɛ^ -acetyltransferase (HsSSAT2) in complex with acetyl coenzyme A. Proteins 64:288–293.

Hanfrey, C., Sommer, S., Mayer, M. J., Burtin, D., and Michael, A. J. (2001). *Arabidopsis* polyamine biosynthesis: absence of ornithine decarboxylase and the mechanism of arginine decarboxylase activity. The Plant Journal 27:551–560.

Hasan, Md. M., Alabdallah, N. M., Alharbi, B. M., Waseem, M., Yao, G., Liu, X.-D., Abd El-Gawad, H. G., El-Yazied, A. A., Ibrahim, M. F. M., Jahan, M. S., et al. (2021). GABA: A Key Player in Drought Stress Resistance in Plants. IJMS 22:10136.

Hegde, S. S., Chandler, J., Vetting, M. W., Yu, M., and Blanchard, J. S. (2007). Mechanistic and Structural Analysis of Human Spermidine/Spermine *N* ^1^ -Acetyltransferase,. Biochemistry 46:7187–7195.

Hennion, F., Bouchereau, A., Gauthier, C., Hermant, M., Vernon, P., and Prinzing, A. (2012). Variation in amine composition in plant species:How it integrates macroevolutionary and environmental signals. American J of Botany 99:36–45.

Jammes, F., Leonhardt, N., Tran, D., Bousserouel, H., Véry, A., Renou, J., Vavasseur, A., Kwak, J. M., Sentenac, H., Bouteau, F., et al. (2014). Acetylated 1,3-diaminopropane antagonizes abscisic acid-mediated stomatal closing in A rabidopsis. The Plant Journal 79:322–333.

Jang, S. J., Wi, S. J., Choi, Y. J., An, G., and Park, K. Y. (2012). Increased Polyamine Biosynthesis Enhances Stress Tolerance by Preventing the Accumulation of Reactive Oxygen Species: T-DNA Mutational Analysis of Oryza sativa Lysine Decarboxylase-like Protein 1. Molecules and Cells 34:251–262.

Kamada-Nobusada, T., Hayashi, M., Fukazawa, M., Sakakibara, H., and Nishimura, M. (2008). A Putative Peroxisomal Polyamine Oxidase, AtPAO4, is Involved in Polyamine Catabolism in Arabidopsis thaliana. Plant and Cell Physiology 49:1272–1282.

Kermode, A. R. (2005). Role of Abscisic Acid in Seed Dormancy. J Plant Growth Regul 24:319–344.

Kilian, J., Whitehead, D., Horak, J., Wanke, D., Weinl, S., Batistic, O., D’Angelo, C., Bornberg-Bauer, E., Kudla, J., and Harter, K. (2007). The AtGenExpress global stress expression data set: protocols, evaluation and model data analysis of UV-B light, drought and cold stress responses. The Plant Journal 50:347–363.

Kim, D. W., Watanabe, K., Murayama, C., Izawa, S., Niitsu, M., Michael, A. J., Berberich, T., and Kusano, T. (2014). Polyamine Oxidase5 Regulates Arabidopsis Growth through Thermospermine Oxidase Activity. Plant Physiology 165:1575–1590.

Kusano, M., Fukushima, A., Arita, M., Jonsson, P., Moritz, T., Kobayashi, M., Hayashi, N., Tohge, T., and Saito, K. (2007). Unbiased characterization of genotype-dependent metabolic regulations by metabolomic approach in Arabidopsis thaliana. BMC Syst Biol 1:53.

Kusano, T., Berberich, T., Tateda, C., and Takahashi, Y. (2008). Polyamines: essential factors for growth and survival. Planta 228:367–381.

Legrandp, Soleilproxima1, Aishima, J., and CV-GPhL (2019). legrandp/xdsme: March 2019 version working with the latest XDS version (Jan 26, 2018) Advance Access published March 28, 2019, doi:10.5281/ZENODO.837885.

Li, L., Dou, N., Zhang, H., and Wu, C. (2021). The versatile GABA in plants. Plant Signaling & Behavior 16:1862565.

Liu, S., Jiang, J., Ma, Z., Xiao, M., Yang, L., Tian, B., Yu, Y., Bi, C., Fang, A., and Yang, Y. (2022). The Role of Hydroxycinnamic Acid Amide Pathway in Plant Immunity. Front. Plant Sci. 13:922119.

Lou, Y.-R., Bor, M., Yan, J., Preuss, A. S., and Jander, G. (2016). Arabidopsis NATA1 acetylates putrescine and decreases defense-related hydrogen peroxide accumulation. Plant Physiol. Advance Access published April 25, 2016, doi:10.1104/pp.16.00446.

Macoy, D. M., Kim, W.-Y., Lee, S. Y., and Kim, M. G. (2015). Biotic stress related functions of hydroxycinnamic acid amide in plants. J. Plant Biol. 58:156–163.

Mattioli, R., Pascarella, G., D’Incà, R., Cona, A., Angelini, R., Morea, V., and Tavladoraki, P. (2022). Arabidopsis N-acetyltransferase activity 2 preferentially acetylates 1,3-diaminopropane and thialysine. Plant Physiology and Biochemistry 170:123–132.

Mesnard, F., Azaroual, N., Marty, D., Fliniaux, M.-A., Robins, R. J., Vermeersch, G., and Monti, J.-P. (2000). Use of 15N reverse gradient two-dimensional nuclear magnetic resonance spectroscopy to follow metabolic activity in Nicotiana plumbaginifolia cell-suspension cultures. Planta 210:446–453.

Montemayor, E. J., and Hoffman, D. W. (2008). The Crystal Structure of Spermidine/Spermine *N* ^1^ -Acetyltransferase in Complex with Spermine Provides Insights into Substrate Binding and Catalysis,. Biochemistry 47:9145–9153.

Moschou, P. N., Wu, J., Cona, A., Tavladoraki, P., Angelini, R., and Roubelakis-Angelakis, K. A. (2012). The polyamines and their catabolic products are significant players in the turnover of nitrogenous molecules in plants. Journal of Experimental Botany 63:5003–5015.

Müller, M., and Munné-Bosch, S. (2015). Ethylene Response Factors: A Key Regulatory Hub in Hormone and Stress Signaling. Plant Physiol. 169:32–41.

Nahar, K., Hasanuzzaman, M., Rahman, A., Alam, Md. M., Mahmud, J.-A., Suzuki, T., and Fujita, M. (2016). Polyamines Confer Salt Tolerance in Mung Bean (Vigna radiata L.) by Reducing Sodium Uptake, Improving Nutrient Homeostasis, Antioxidant Defense, and Methylglyoxal Detoxification Systems. Front. Plant Sci. 7.

Nambara, E. (2017). Abscisic Acid. In Encyclopedia of Applied Plant Sciences, pp. 361–366. Elsevier.

Pegg, A. E. (2008). Spermidine/spermine-*N* ^1^ -acetyltransferase: a key metabolic regulator. American Journal of Physiology-Endocrinology and Metabolism 294:E995–E1010.

Priya, M., Sharma, L., Kaur, R., Bindumadhava, H., Nair, R. M., Siddique, K. H. M., and Nayyar, H. (2019). GABA (γ-aminobutyric acid), as a thermo-protectant, to improve the reproductive function of heat-stressed mungbean plants. Sci Rep 9:7788.

Qiu, X.-M., Sun, Y.-Y., Ye, X.-Y., and Li, Z.-G. (2020). Signaling Role of Glutamate in Plants. Front. Plant Sci. 10:1743.

Regla-Márquez, C. F., Canto-Flick, A., Avilés-Viñas, S. A., Valle-Gough, R. E., Pérez-Pastrana, J., García-Villalobos, F. J., and Santana-Buzzy, N. (2016). Cadaverine: a common polyamine in zygotic embryos and somatic embryos of the species Capsicum chinense Jacq. Plant Cell Tiss Organ Cult 124:253–264.

Rhee, H. J., Kim, E., and Lee, J. K. (2007). Physiological polyamines: simple primordial stress molecules. J Cellular Molecular Medi 11:685–703.

Rossi, F., Rizzotti, L., Felis, G. E., and Torriani, S. (2014). Horizontal gene transfer among microorganisms in food: Current knowledge and future perspectives. Food Microbiology 42:232–243.

Rosso, M. G., Li, Y., Strizhov, N., Reiss, B., Dekker, K., and Weisshaar, B. (2003). An Arabidopsis thaliana T-DNA mutagenized population (GABI-Kat) for flanking sequence tag-based reverse genetics. Plant Mol Biol 53:247–259.

Sagor, G. H. M., Inoue, M., Kim, D. W., Kojima, S., Niitsu, M., Berberich, T., and Kusano, T. (2015). The polyamine oxidase from lycophyte *Selaginella lepidophylla* (SelPAO5), unlike that of angiosperms, back-converts thermospermine to norspermidine. FEBS Letters 589:3071–3078.

Schmid, M., Davison, T. S., Henz, S. R., Pape, U. J., Demar, M., Vingron, M., Schölkopf, B., Weigel, D., and Lohmann, J. U. (2005). A gene expression map of Arabidopsis thaliana development. Nat Genet 37:501–506.

Sen, S., Ghosh, D., and Mohapatra, S. (2018). Modulation of polyamine biosynthesis in Arabidopsis thaliana by a drought mitigating Pseudomonas putida strain. Plant Physiology and Biochemistry 129:180–188.

Shahul Hameed, U., Haider, I., Jamil, M., Guo, X., Zarban, R. A., Kim, D., Al-Babili, S., and Arold, S. T. (2022). Structural basis for specific inhibition of the highly sensitive ShHTL7 receptor. EMBO Reports 23:e54145.

Shen, X., Li, Y., Pan, Y., and Zhong, S. (2016). Activation of HLS1 by Mechanical Stress via Ethylene-Stabilized EIN3 Is Crucial for Seedling Soil Emergence. Front. Plant Sci. 7.

Sita, K., and Kumar, V. (2020). Role of Gamma Amino Butyric Acid (GABA) against abiotic stress tolerance in legumes: a review. Plant Physiol. Rep. 25:654–663.

Sobieszczuk-Nowicka, E. (2017). Polyamine catabolism adds fuel to leaf senescence. Amino Acids 49:49–56.

Suza, W. P., and Staswick, P. E. (2008). The role of JAR1 in Jasmonoyl-l-isoleucine production during Arabidopsis wound response. Planta 227:1221–1232.

Takahashi, T., and Kakehi, J.-I. (2010). Polyamines: ubiquitous polycations with unique roles in growth and stress responses. Annals of Botany 105:1–6.

Takahashi, Y., Tahara, M., Yamada, Y., Mitsudomi, Y., and Koga, K. (2018). Characterization of the Polyamine Biosynthetic Pathways and Salt Stress Response in Brachypodium distachyon. J Plant Growth Regul 37:625–634.

Tavladoraki, P., Cona, A., Federico, R., Tempera, G., Viceconte, N., Saccoccio, S., Battaglia, V., Toninello, A., and Agostinelli, E. (2012). Polyamine catabolism: target for antiproliferative therapies in animals and stress tolerance strategies in plants. Amino Acids 42:411–426.

Tavladoraki, P., Cona, A., and Angelini, R. (2016). Copper-Containing Amine Oxidases and FAD-Dependent Polyamine Oxidases Are Key Players in Plant Tissue Differentiation and Organ Development. Front. Plant Sci. 7.

Thurow, C., Krischke, M., Mueller, M. J., and Gatz, C. (2020). Induction of Jasmonoyl-Isoleucine (JA-Ile)-Dependent JASMONATE ZIM-DOMAIN (JAZ) Genes in NaCl-Treated Arabidopsis thaliana Roots Can Occur at Very Low JA-Ile Levels and in the Absence of the JA/JA-Ile Transporter JAT1/AtABCG16. Plants 9:1635.

Tzin, V., Fernandez-Pozo, N., Richter, A., Schmelz, E. A., Schoettner, M., Schäfer, M., Ahern, K. R., Meihls, L. N., Kaur, H., Huffaker, A., et al. (2015). Dynamic maize responses to aphid feeding are revealed by a time series of transcriptomic and metabolomic assays. Plant Physiol. Advance Access published September 16, 2015, doi:10.1104/pp.15.01039.

Vonrhein, C., Flensburg, C., Keller, P., Sharff, A., Smart, O., Paciorek, W., Womack, T., and Bricogne, G. (2011). Data processing and analysis with the *autoPROC* toolbox. Acta Crystallogr D Biol Crystallogr 67:293–302.

Waese, J., Fan, J., Pasha, A., Yu, H., Fucile, G., Shi, R., Cumming, M., Kelley, L. A., Sternberg, M. J., Krishnakumar, V., et al. (2017). ePlant: Visualizing and Exploring Multiple Levels of Data for Hypothesis Generation in Plant Biology. Plant Cell 29:1806– 1821.

Wimalasekera, R., Tebartz, F., and Scherer, G. F. E. (2011). Polyamines, polyamine oxidases and nitric oxide in development, abiotic and biotic stresses. Plant Science 181:593–603.

Xia, J., and Wishart, D. S. (2016). Using MetaboAnalyst 3.0 for Comprehensive Metabolomics Data Analysis. CP in Bioinformatics 55.

Yoda, H., Hiroi, Y., and Sano, H. (2006). Polyamine Oxidase Is One of the Key Elements for Oxidative Burst to Induce Programmed Cell Death in Tobacco Cultured Cells. Plant Physiology 142:193–206.

Yoda, H., Fujimura, K., Takahashi, H., Munemura, I., Uchimiya, H., and Sano, H. (2009). Polyamines as a common source of hydrogen peroxide in host- and nonhost hypersensitive response during pathogen infection. Plant Mol Biol 70:103–112.

Zahedi, K., Barone, S., and Soleimani, M. (2022). Polyamines and Their Metabolism: From the Maintenance of Physiological Homeostasis to the Mediation of Disease. Medical Sciences 10:38.

