## Supplementary_figures_tables for "Regulation of Arabidopsis polyamine acetylation by NATA1 and NATA2"

### Supplementary files

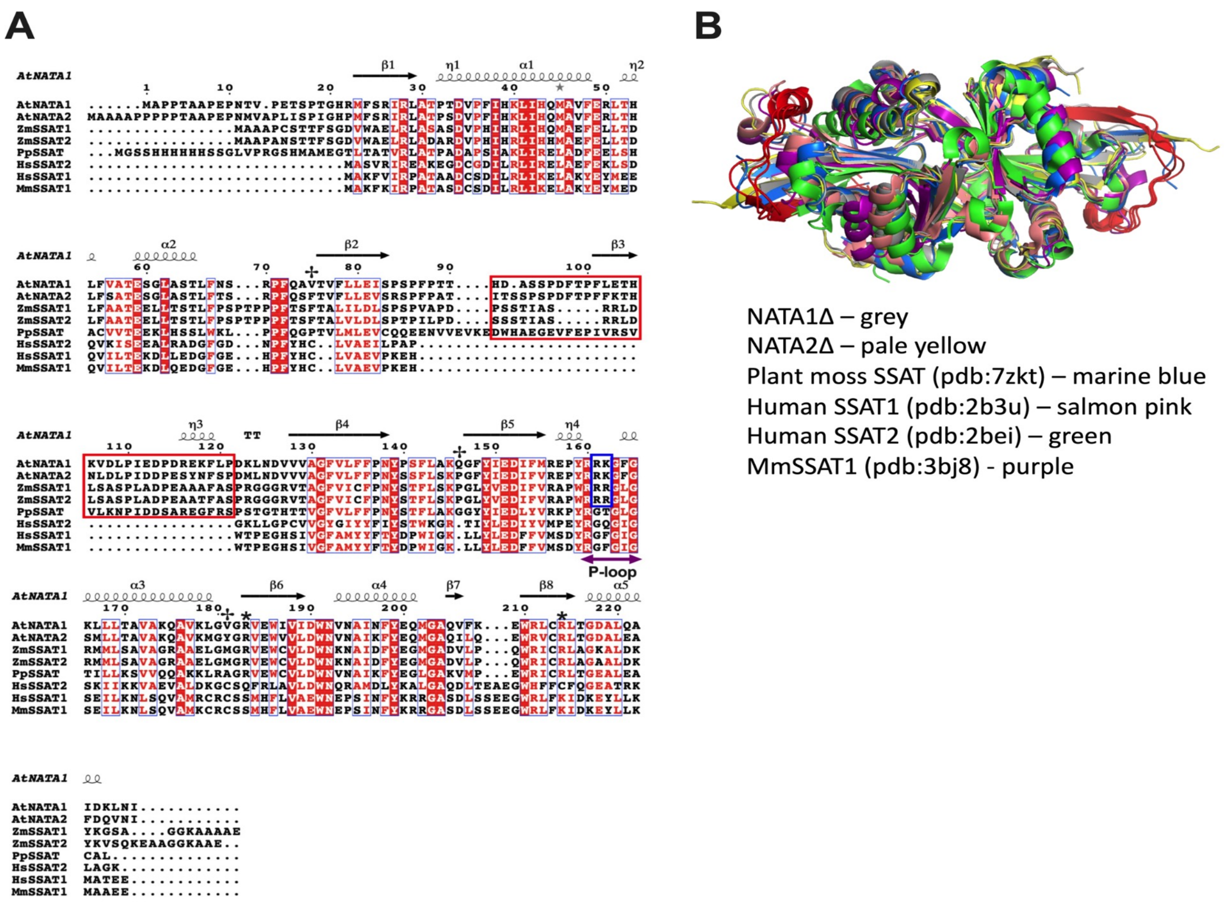


**Supplementary Figure S1. Comparison of SSAT homologues. A**) Multiple sequence alignment of the SSATs from different species. The plant SSAT–specific loop sequences are boxed in red. The additional basic residues uniquely present in the P-loops of plant SSATs are boxed in blue. The two arginines from the di-arginine motif in the catalytic pocket are marked by an asterisk (*), and the three residue positions important for the heat stability of NATA2 versus NATA2 are marked by a cross (✢). Secondary structure derived from NATA1 is shown above the sequences. **B**) Superimposition of the PDB structures of SSAT dimers. NATA1Δ - grey; NATA2Δ - yellow; *Pp*SSAT (PDB 7ZHC) - blue; *Hs*SSAT1 (PDB 2B3U) - salmon; *Hs*SSAT2 (PDB 2BEI) - green; *Mm*SSAT1 (PDB 3BJ8) - purple. The additional loop present in plant SSATs is shown in red.

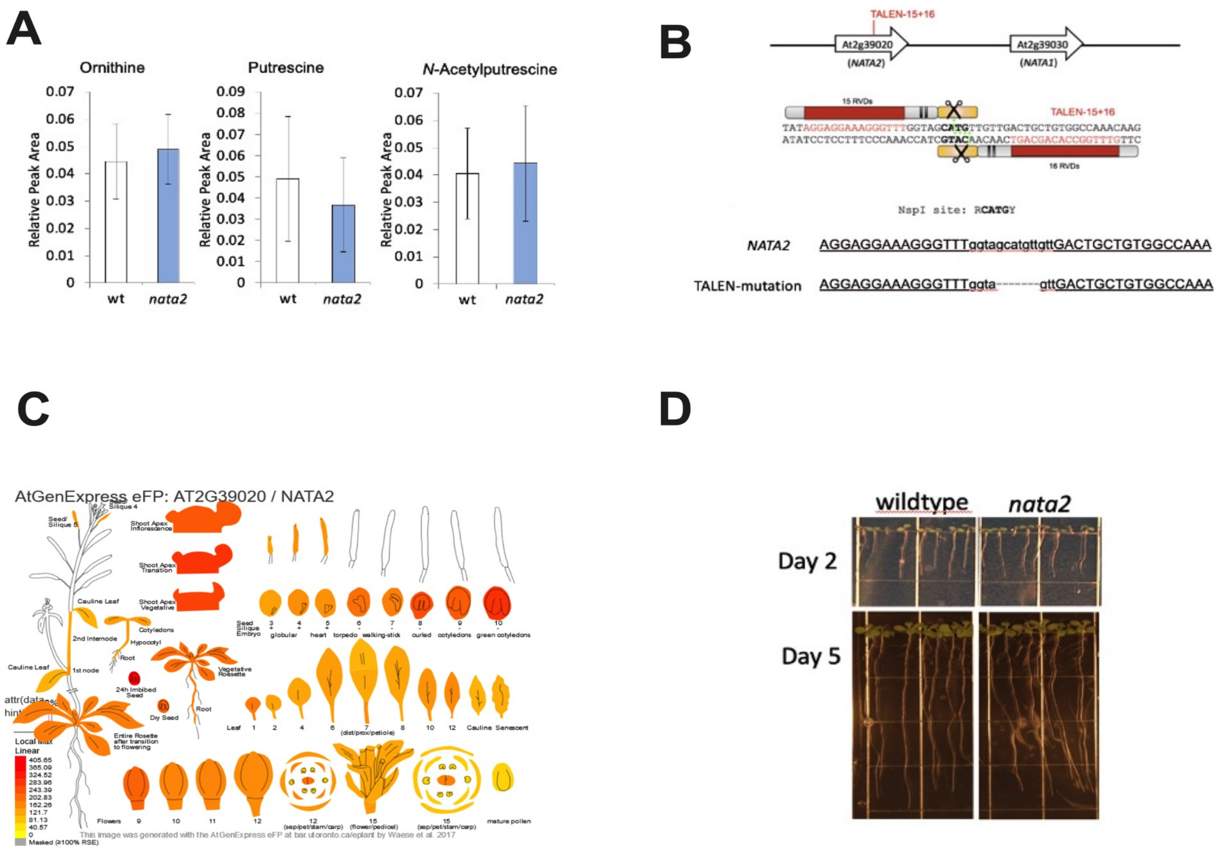


**Supplementary Figure S2. Design and effects of nata2 mutant plants. A**) Accumulation of the polyamines ornithine (*left*), putrescine (*middle*) and *N*-acetylputrescine (*right*) in rosette leaves of wild-type *A. thaliana* and *nata2* mutant plants, as measured by HPLC. Mean ± SE of N=9. **B**) NATA2 sequence changes due to the TALENS. (*Top*) Schematic overview of the strategy to disrupt *NATA2* in the *nata1* mutant background. RVD, Repeat Variable Diresidue. (*Bottom*) Sanger sequencing result of the selected line used for generating a *nata1 nata2* double mutant. **C**) Spatial and temporal expression of *NATA2,* At2g39020. Data are from ePlant (https://bar.utoronto.ca/eplant/; Waese et al. 2017). **D**) *A. thaliana nata2* seedlings show no obvious variation in the growth phenotype compared to Col-0 wild-type seedlings after two and five days growth on half-strength Murashige and Skoog agar plates.

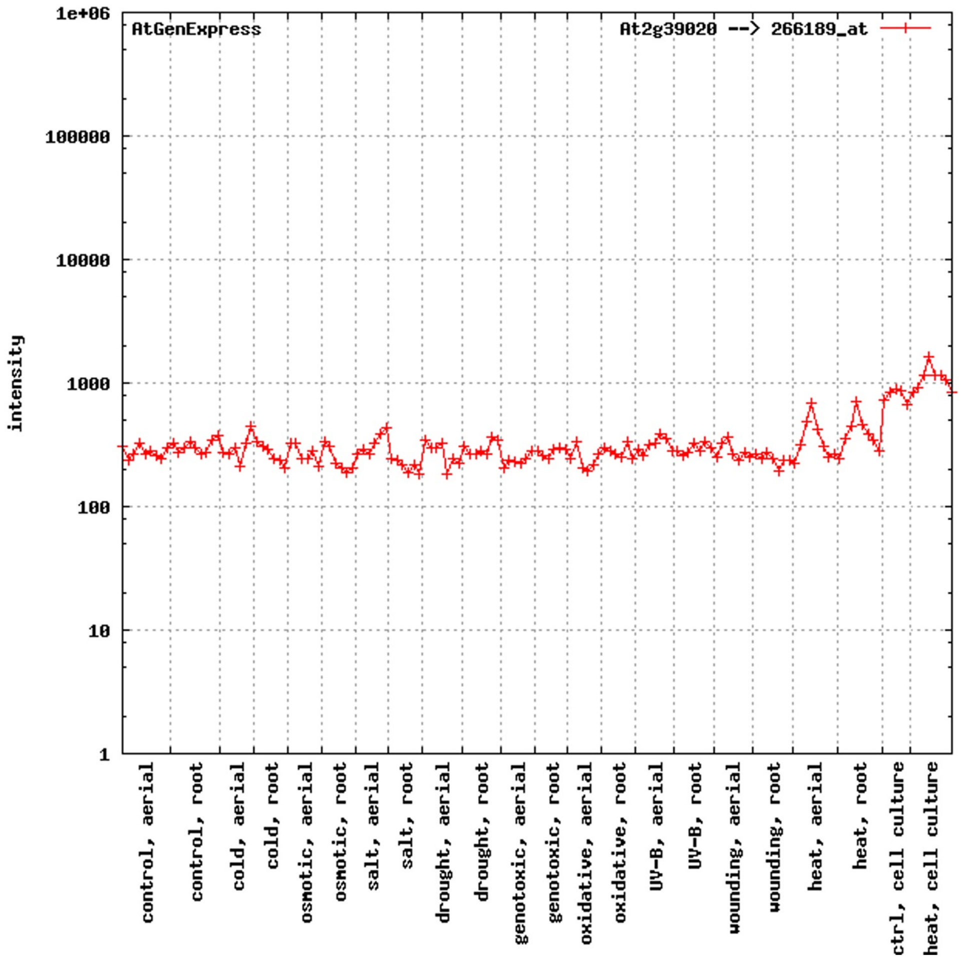


**Supplementary Figure S3.** ***NATA2*** **gene expression in response to abiotic stress. http://jsp.weigelworld.org/expviz/; (Schmid et al., 2005).**

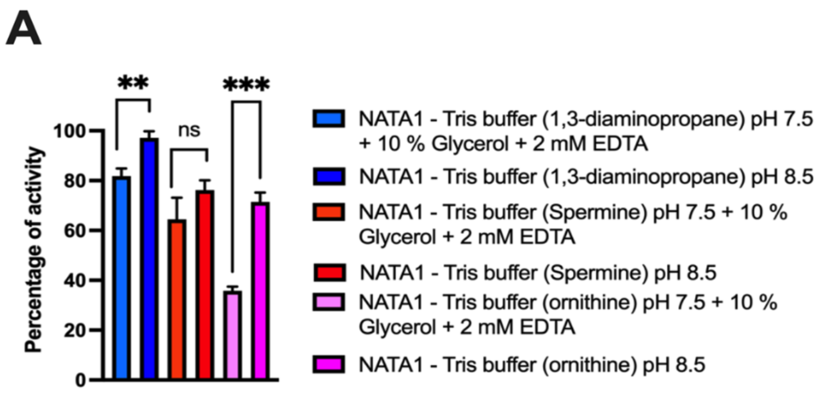


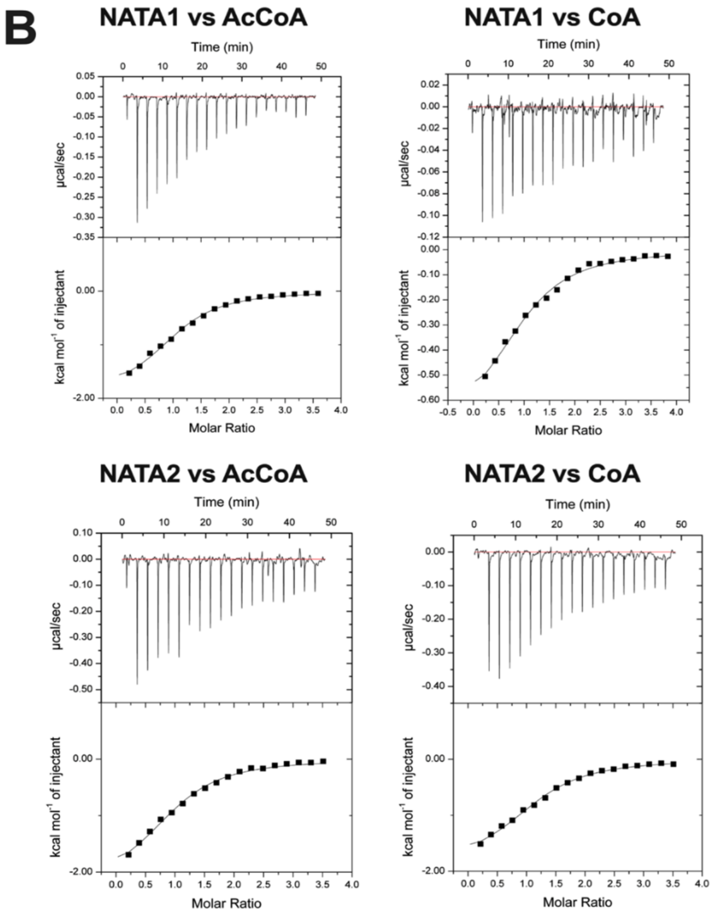


C

| cell | syringe | N | $K_d$ (µM) | ΔG (kJ/mol) | ΔH (kJ/mol) | TΔS (kJ/mol/K) |
|---|---|---|---|---|---|---|
| NATA1Δ | acCoA | 1.04 ± 0.10 | 9.34 ± 1.52 | -28.75 ± 2.95 | -8.04 ± 0.95 | 20.71 ± 2.55 |
| NATA1Δ | CoA | 0.95 ± 0.15 | 11.05 ± 1.21 | -28.36 ± 2.50 | -3.05 ± 0.27 | 25.31 ± 2.25 |
| NATA2Δ | acCoA | 0.98 ± 0.10 | 8.62 ± 1.05 | -28.86 ± 1.95 | -9.41 ± 0.85 | 19.45 ± 1.55 |
| NATA2Δ | CoA | 1.15 ± 0.15 | 7.77 ± 0.25 | -29.12 ± 2.15 | -7.91 ± 0.65 | 21.21 ± 2.25 |

**Supplementary Figure S4. NATA1Δ catalytic activity in the presence of additives (ac)CoA binding. A)** Comparison of the catalytic activity of NATA1Δ against its substrates in the presence of ethylene glycol, glycerol, and hexylene glycol in buffer Tris8.5 and Tris7.5 at 25 ˚C. **B**) ITC investigating binding of (ac)CoA to NATA1Δor NATA2Δ. *Top panels*: raw data showing heats for each injection. *Bottom panels*: Integrated heats from top panel. **C**) Table of binding affinities and the thermodynamic parameters calculated from (B).

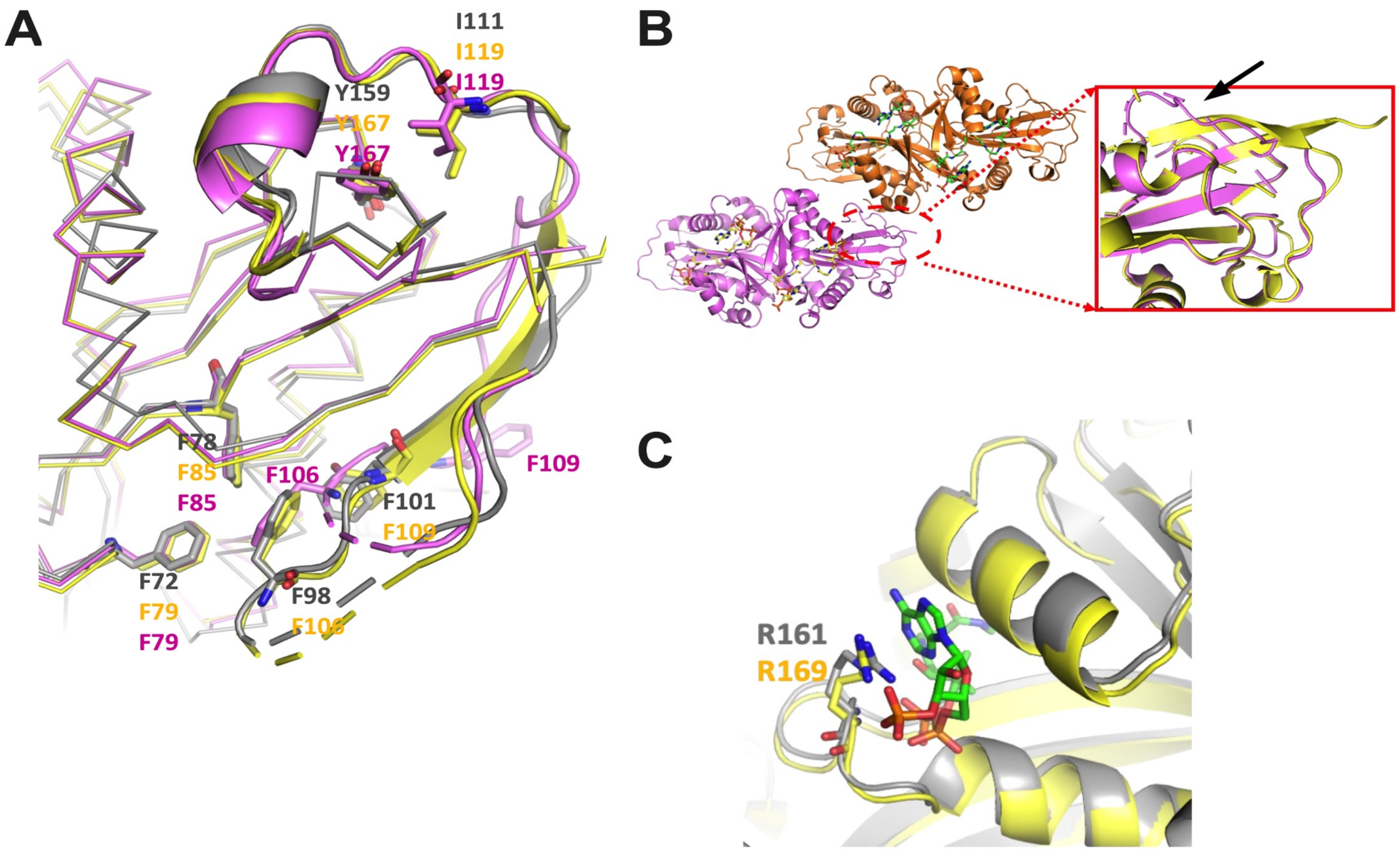


**Supplementary Figure S5. Structural details of NATA1Δ/NATA2Δ.** A) Zoom view into the plants-specific insert and its interactions with the conserved NATA core. Shown are superimposed structures of NATA1Δ bound to HEPES and CoA (grey), NATA2Δ bound to HEPES and acCoA (yellow) and NATA2Δ bound with di-CoA (magenta), produced in-cristallo. Key residues involved in the interaction between the loop and core residues are shown as sticks. **B**) Structural rearrangement of the β-strand from the plant-specific insert caused by a symmetry-related molecule (orange) in the NATA2Δ bound to di-CoA crystal structure. The inlay zooms into the displaced β-strand of the di-CoA–bound NATA2Δ structure (marked by an arrow). Superimposed is the structure of NATA2Δ bound to HEPES-CoA (yellow) with the unaffected β-strand conformation. **C**) Position of the additional arginine present in the P-loop of higher plants, including Arabidopsis. Arginines are shown as stick models (NATA1Δ: grey carbons; NATA2Δ: yellow carbons). The acCoA is shown as a stick model with green carbons.

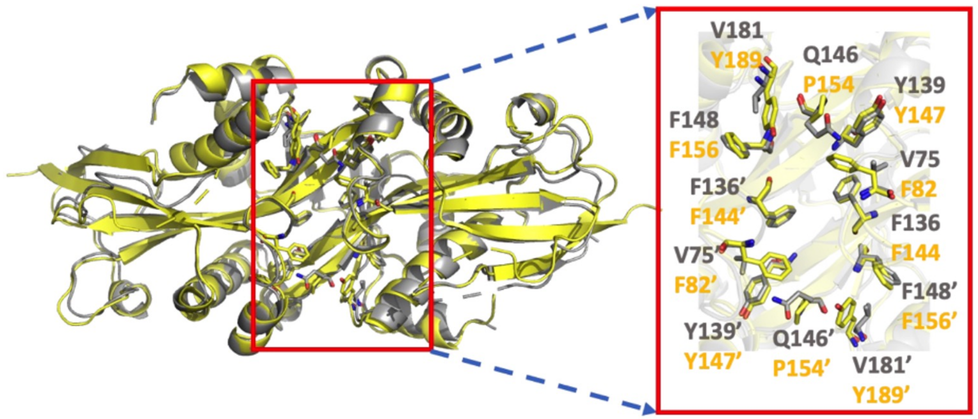


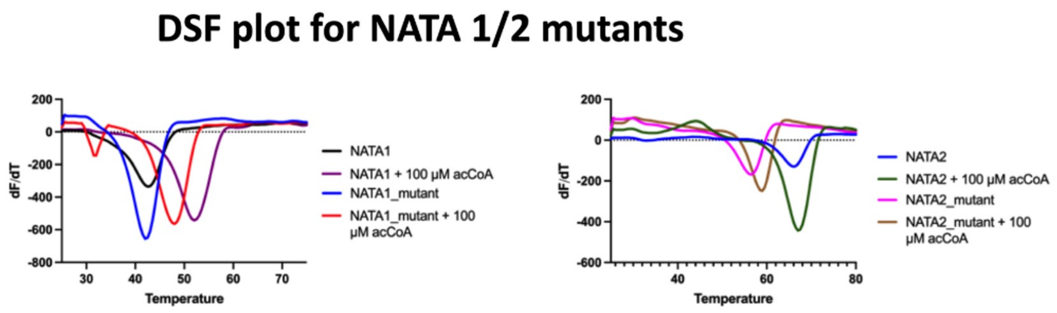


**B**

| A | B | C | D | E | F | G | H | I | J | K | L | M | N | O | P | Q | R | S | T | U | V | W | X |
|---|---|---|---|---|---|---|---|---|---|---|---|---|---|---|---|---|---|---|---|---|---|---|---|
| Pdb | Total Energy | Backbone Hb | Sidechain Hb | Van der Waal | Electrostatics | Solvation Pola | Solvation Hyd | Van der Waal | Entropy Side | Entropy Main | Sloop Entropy | Mloop Entrop | Cis Bond | Torsional Clas | Backbone Cla | Helix Dipole | Water Bridge | Disulfide | Electrostatic | Partial Covale | Energy Ionisa | Entropy Comp | Residue Number |
| ./NATA1_dimer_apo_Repair.pdb | -75.6265 | -281.229 | -101.642 | -495.615 | -26.1436 | 612.717 | -678.022 | 10.2772 | 258.95 | 617.403 | 0 | 0 | 2.24811 | 8.19132 | 221.315 | -2.45097 | 0 | 0 | -1.24466 | 0 | 0.933285 | 0 | 414 |
| ./NATA1mut_NATA2_Repair.pdb | -97.946 | -283.034 | -103.597 | -501.066 | -28.116 | 615.284 | -685.763 | 13.1863 | 261.116 | 610.297 | 0 | 0 | 2.24811 | 4.66311 | 216.514 | -2.68357 | 0 | 0 | -1.41287 | 0 | 0.93171 | 0 | 414 |
| ./NATA2_dimer_apo_Repair.pdb | -98.9751 | -290.835 | -111.416 | -498.333 | -24.5159 | 613.759 | -672.943 | 8.95739 | 259.385 | 615.144 | 0 | 0 | 2.24811 | 3.36522 | 216.871 | -2.54737 | 0 | 0 | -2.11735 | 0 | 0.873165 | 0 | 416 |
| ./NATA2mut_NATA1_Repair.pdb | -85.9567 | -289.45 | -108.902 | -494.333 | -24.412 | 614.506 | -666.809 | 8.14063 | 259.513 | 613.905 | 0 | 0 | 2.24811 | 3.45052 | 220.664 | -2.53527 | 0 | 0 | -2.12695 | 0 | 0.849409 | 0 | 416 |

**Supplementary Figure S6. Molecular basis of NATA2 heat stability and its validation by mutations. A**) Superimposition of NATA1Δ bound to HEPES and CoA (grey) and NATA2Δ bound to HEPES and acCoA (yellow). Zoom figure shows the central core residues identified as modulating the stability of the proteins. **B**) Table showing the heat stability values of NATA1Δ and NATA2Δ and its mutants using the FoldX server. All values are in kcal/mol. **C**) DSF traces showing the heat stability of NATA1Δ and NATA2Δ and mutants.

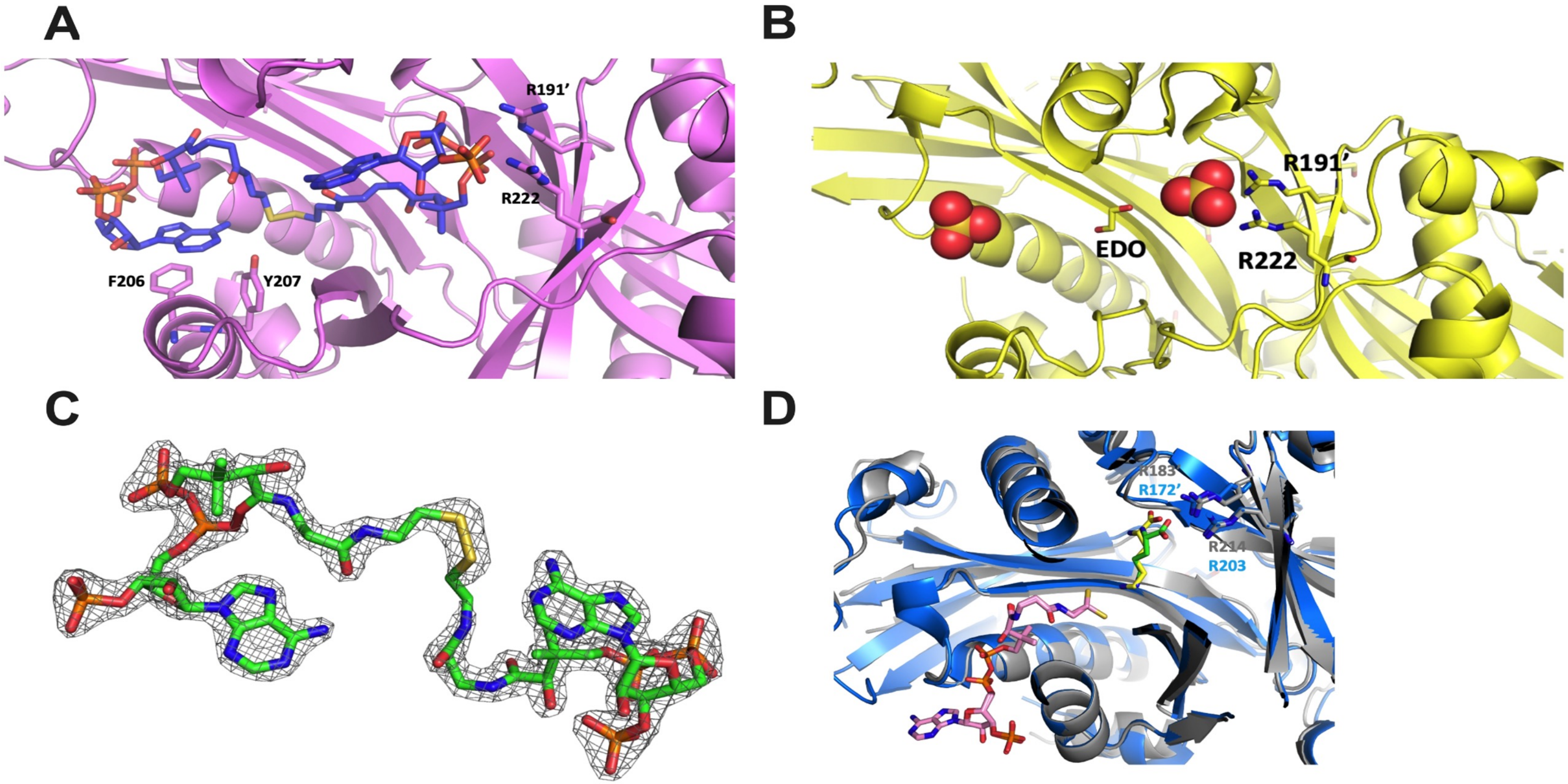


**Supplementary Figure S7. Details of NATA1Δ, NATA2 and NATA2Δ and their ligand interactions. A**) Zoom view of NATA2Δ (magenta) bound to di-CoA (blue carbons). di-CoA and NATA2Δ residues involved in binding are shown as sticks. **B**) Zoom view of the structure of apo-NATA2 (yellow) crystallised from a solution containing full-length protein. Sulphate atoms shown as sphere models and the NATA2 residues are shown as sticks. The ethanediol (EDO) bound in the catalytic pocket is shown as a stick model with carbons in yellow. **C**) 2Fo-Fc omit map of di-CoA contoured at 1**σ**. **D**) Superimposition of NATA1Δ bound to ornithine and CoA (grey), and of *Pp*SSAT bound to lysine and acCoA (pdb: 7zkt; blue). Stick models show ornithine (green carbons), lysine (yellow), acCoA (pink) and the di-arginines involved in binding.

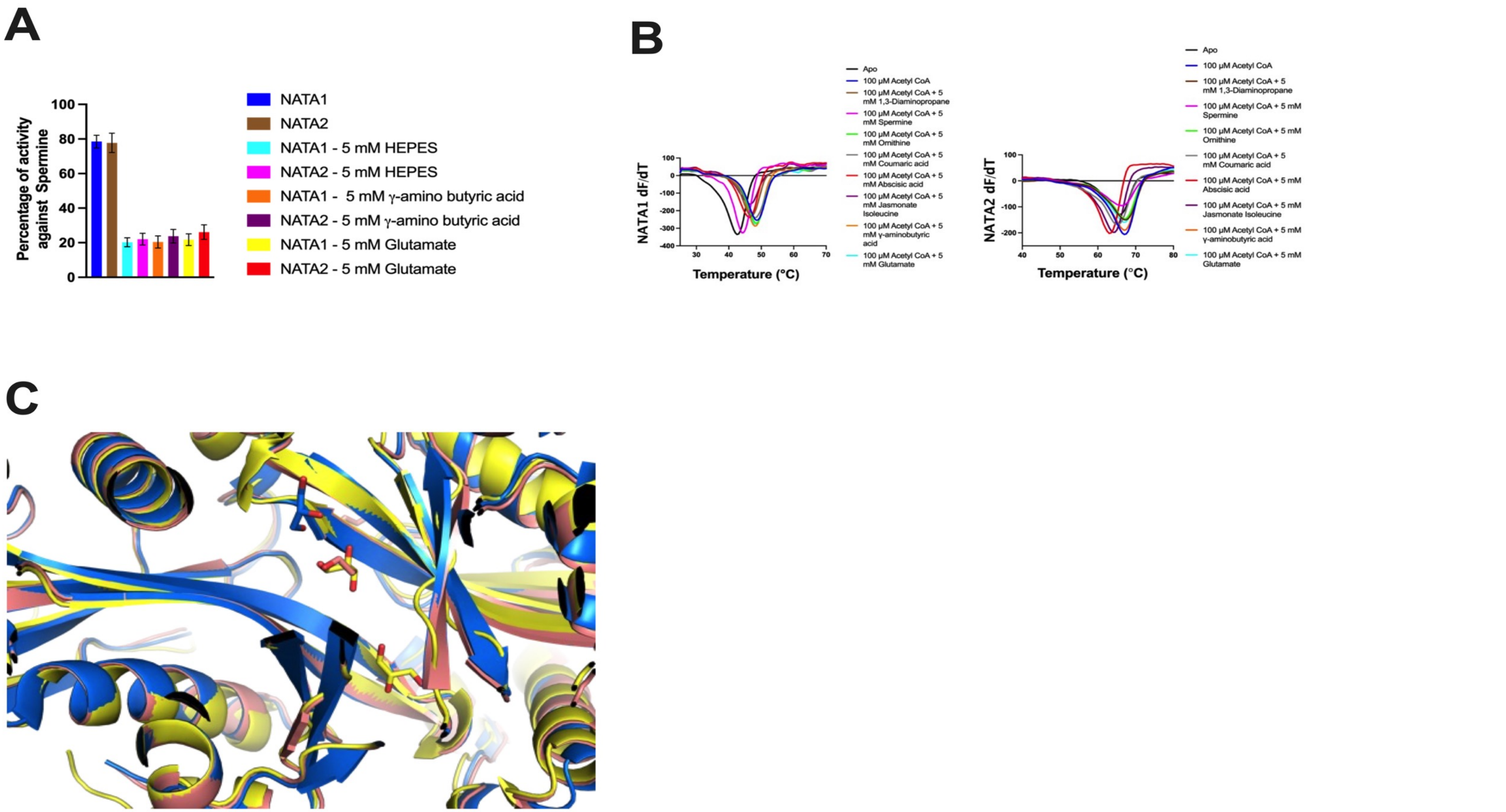


**Supplementary Figure S8. Additional experimental and structural data. A**) Catalytic activity of NATA2Δ on spermine in the presence of acidic compounds. **B**) DSF data indicating heat stability of NATA1Δ and NATA2Δ in the presence of acidic endogenous metabolites. **C**) Glycerol and ethanediol bound in the catalytic pocket are shown as sticks, colour-coded according to the crystal structure in which they were identified: NATA2Δ bound to HEPES and CoA (yellow), full-length apo-NATA2 (orange), and *Pp*SSAT (blue; pdb 7zkt).

| Protein | PDB | Resolution (Å) | Organism | additional compounds | Conformation (open/closed) | Reference |
|---|---|---|---|---|---|---|
| NATA1Δ | 8XBN | 1.35 | A. thaliana | CoA, HEPES | closed | This work |
| NATA1Δ | 8XJ5 | 2.04 | A. thaliana | CoA, Ornithine | closed | This work |
| NATA1Δ | 8XBP | 1.99 | A. thaliana | acCoA, calcium | closed | This work |
| NATA2 | 8RMZ | 1.74 | A. thaliana | Ethanediol, sulfate | open | This work |
| NATA2Δ | 8XJ9 | 1.10 | A. thaliana | PEG | open | This work |
| NATA2Δ | 8XJB | 1.45 | A. thaliana | Spermine | open | This work |
| NATA2Δ | 8XJF | 2.00 | A. thaliana | acCoA, HEPES, **γ**-aminobutyric acid, glycerol | closed | This work |
| NATA2Δ | 8XJH | 1.75 | A. thaliana | di-CoA | open | This work |
| NATA2 | 7OVV | 1.45 | A. thaliana | di-CoA | open | unpublished |
| PpSSAT | 7ZHC | 1.82 | P. patens | acCoA, PEG, glycerol | open and closed molecules | https://doi.org/10.1111/tpj.16148 |
| PpSSAT | 7ZKT | 2.06 | P. patens | CoA, lysine, sulfate, sodium, ethanediol | open and closed | https://doi.org/10.1111/tpj.16148 |
| SSAT2 | 2BEI | 1.84 | Human | acCoA | open and closed | https://doi.org/10.1002/prot.20967 |
| SSAT1 | 3BJ7 | 2.20 | Mouse | CoA | open and closed | https://doi.org/10.1021/bi8009357 |
| SSAT1 | 3BJ8 | 2.30 | Mouse | CoA, Spermine | open and closed | https://doi.org/10.1021/bi8009357 |
| SSAT1 | 2B4B | 2.00 | Human | COA, BE333 | open and closed | https://doi.org/10.1073/pnas.0511008103 |

| | | | | | | |
|---|---|---|---|---|---|---|
| SSAT1 | 2B4D | 2.00 | Human | CoA | closed | https://doi.org/10.1073/pnas.0511008103 |
| SSAT1 | 2F5I | 2.30 | Human | | open and closed | https://doi.org/10.1002/prot.20965 |
| SSAT1 | 2JEV | 2.30 | Human | N1-spermine-acCoA | open and closed | https://doi.org/10.1021/bi700256z |

**Supplementary Table 1. Summary of NATA/SSAT structures.**

| | NATA2 apo | NATA2Δ apo | NATA2Δ + Spermine | NATA2Δ + HEPES + AcCoA | NATA2Δ + di-CoA | NATA1Δ + HEPES + CoA | NATA1Δ + Ornithine + CoA | NATA1Δ + AcCoA |
|---|---|---|---|---|---|---|---|---|
| PDB code | **8RMZ** | **8XJ9** | **8XJB** | **8XJF** | **8XJH** | **8XBN** | **8XJ5** | **8XBP** |
| Space group Cell parameters (Å,°) | $P4_32_12$ 101.2 101.2 48.9 90 90 90 | C121 73.7 54.8 51.7 90 113.90 90 | C121 73.7 54.8 51.9 90 113.6 90 | C2221 60.0 69.7 104.3 90 90 90 | P212121 55.4 85.5 86.1 90 90 90 | P21212 64.2 76.6 82.08 90 90 90 | P212121 46 74 110.3 90 90 90 | C121 148.3 50.8 60.2 90 92.33 90 |
| Resolution (Å) | 71.58 -1.74 (1.84-1.74) | 36.61 - 1.10 (1.13 - 1.10) | 33.77 - 1.45 (1.50 - 1.45) | 45.51 - 2.00 (2.07 - 2.00) | 46.63 - 1.75 (1.81 - 1.75) | 42.24 - 1.35 (1.39 - 1.35) | 42.49 - 2.04 (2.11 - 2.04) | 47.66 - 1.99 (2.066 - 1.99) |
| observed reflections | 570015 (24736) | 508642 (44082) | 227246 (17550) | 82977 (3772) | 597189 (58764) | 1172428 (102111) | 327707 (30111) | 181604 (7554) |
| No. of unique reflections | 21764 (1088) | 75933 (7263) | 33619 (3140) | 15067 (1088) | 41905 (4113) | 88772 (8164) | 24570 (2288) | 28289 (1699) |
| Completeness (%) | 81.6 (26.7) | 98.2 (93.4) | 99.4 (93.8) | 98.1 (77.1) | 99.9 (99.7) | 98.3 (91.3) | 99.4 (94.1) | 91.8 (55.3) |
| $R_{merge}$ (%) | 10.7 (312.3) | 9.30 (179.6) | 6.14 (123.2) | 7.17 (287.1) | 10.17 (210.5) | 5.93 (218.5) | 18.42 (242.6) | 27.52 (246.8) |
| $R_{pim}$ (%) | 2.1 (66.7) | 3.83 (77.6) | 2.53 (55.8) | 3.28 (141) | 2.79 (57.1) | 1.72 (63.0) | 5.23 (68.1) | 11.59 (129) |
| I/s (I) | 14.1 (1.4) | 8.53 (0.9) | 13.91 (1) | 6.20 (0.7) | 13.46 (0.8) | 19.54 (1) | 8.13 (0.9) | 7.44 (0.5) |
| $CC_{1/2}$ | 0.99 (0.56) | 0.99 (0.41) | 0.99 (0.47) | 0.99 (0.35) | 0.99 (0.51) | 0.99 (0.41) | 0.99 (0.49) | 0.99 (0.30) |
| $R_{cryst}$ (%) | 20.76 | 17.05 | 16.42 | 23.48 | 21.26 | 16.24 | 19.99 | 25.21 |
| $R_{free}$ (%) | 22.56 | 19.90 | 19.80 | 26.92 | 24.79 | 20.19 | 25.10 | 28.82 |
| rms bond (Å) | 0.009 | 0.02 | 0.01 | 0.01 | 0.01 | 0.02 | 0.01 | 0.01 |

| | | | | | | | | |
|---|---|---|---|---|---|---|---|---|
| rms angle (°) | 1.02 | 1.76 | 1.39 | 1.58 | 1.69 | 1.70 | 1.27 | 1.54 |
| Average B (Å$^2$)<br>Protein<br>Ligand<br>Solvent | 47.0<br>58.5<br>51.3 | 21.62<br>20.86<br>34.14<br>29.64 | 29.83<br>28.64<br>54.78<br>40.95 | 35.79<br>34.47<br>59.62<br>44.37 | 31.65<br>31.35<br>31.74<br>42.16 | 31.06<br>30.19<br>43.84<br>35.68 | 36.03<br>35.43<br>38.29<br>51.16 | 25.36<br>23.07<br>68.22<br>44.09 |
| [a] Clashscore<br>MolProbity score | 1.46<br>0.96 | 0.89<br>0.77 | 0.9<br>0.77 | 2.63<br>1.15 | 3.28<br>1.23 | 4.91<br>1.40 | 3.84<br>1.61 | 5.72<br>1.90 |
| [a] Ramachandran plot (%)<br>Favoured<br>Outliers | 97.62<br>0.48 | 99.00<br>1.00 | 99.00<br>1.00 | 96.04<br>2.48 | 98.20<br>0.51 | 97.26<br>0.28 | 97.65<br>2.06 | 96.60<br>3.10 |

Values for the highest resolution shell are in parentheses
$CC_{1/2}$ = percentage of correlation between intensities from random hall-dataset
[a]Calculated with MolProbity

**Supplementary Table 2. Crystallographic data.**

#### Supplementary Table 3A

| **Protein** | **Ligand** | **$T_m$ (°C)** |
|---|---|---|
| **NATA1Δ** | apo | 42.5 ± 0.7 |
| **NATA1Δ** | 25 µM acCOA | 46.5 ± 0.5 |
| **NATA1Δ** | 50 µM acCOA | 53 ± 0.3 |
| **NATA1Δ** | 100 µM acCOA | 52.5 ± 0.6 |
| **NATA1Δ** | 100 µM Ornithine | 42.5 ± 0.6 |
| **NATA1Δ** | 100 µM Putrescine | 42.5 ± 0.4 |
| **NATA1Δ** | 100 µM Spermine | 43.2 ± 0.5 |
| **NATA1Δ** | 100 µM 1,3 – Diaminopropane | 42.5 ± 0.7 |
| **NATA2Δ** | apo | 66.5 ± 0.5 |
| **NATA2Δ** | 25 µM acCoA | 68.0 ± 0.5 |
| **NATA2Δ** | 50 µM acCoA | 67.5 ± 0.6 |
| **NATA2Δ** | 100 µM acCoA | 67.5 ± 0.7 |
| **NATA2Δ** | 100 µM Ornithine | 65.2 ± 0.3 |
| **NATA2Δ** | 100 µM Putrescine | 66.0 ± 0.7 |

| | | |
|---|---|---|
| **NATA2Δ** | 100 μM Spermine | 66.0 ± 0.5 |
| **NATA2Δ** | 100 μM 1,3 – Diaminopropane | 66.0 ± 0.8 |
| **NATA1Δ V75F/Q146P/V181Y** | apo | 41.5 ± 0.5 |
| **NATA1Δ V75F/Q146P/V181Y** | 100 μM acCoA | 48.0 ± 0.4 |
| **NATA2Δ F82V/P154Q/Y189V** | apo | 56.5 ± 0.5 |
| **NATA2Δ F82V/P154Q/Y189V** | 100 μM acCoA | 59.0 ± 0.7 |

#### Supplementary Table 3B

| **Protein** | **Ligand** | $T_m$ **(°C)** |
|---|---|---|
| **NATA1Δ** | apo | 42.5 ± 0.7 |
| **NATA1Δ** | 100 μM acCoA | 52.5 ± 0.6 |
| **NATA1Δ** | 100 μM acCoA + 5 mM 1,3 – Diaminopropane | 53 ± 0.3 |
| **NATA1Δ** | 100 μM acCoA + 5 mM Spermine | 47.5 ± 0.6 |
| **NATA1Δ** | 100 μM acCoA + 5 mM Ornithine | 52.0 ± 1 |
| **NATA1Δ** | 100 μM acCoA + 5 mM Coumaric acid | 50.5 ± 0.7 |
| **NATA1Δ** | 100 μM acCoA + 5 mM Abscisic acid | 51.0 ± 0.6 |
| **NATA1Δ** | 100 μM acCoA + Jasmonate isoleucine | 50.5 ± 0.5 |
| **NATA1Δ** | 100 μM acCoA + **γ**-aminobutyric acid | 52.5 ± 0.6 |
| **NATA1Δ** | 100 μM acCoA + Glutamate | 52.3 ± 0.5 |
| **NATA2Δ** | apo | 66.5 ± 0.5 |

| | | |
|---|---|---|
| **NATA2Δ** | 100 µM acCoA | 67.5 ± 0.7 |
| **NATA2Δ** | 100 µM acCoA + 5 mM 1,3 – Diaminopropane | 67.5 ± 0.6 |
| **NATA2Δ** | 100 µM acCoA + 5 mM Spermine | 67.0 ± 0.5 |
| **NATA2Δ** | 100 µM acCoA + 5 mM Ornithine | 67.2 ± 0.3 |
| **NATA2Δ** | 100 µM acCoA + 5 mM Coumaric acid | 66.0 ± 0.7 |
| **NATA2** | 100 µM acCoA + 5 mM Abscisic acid | 63.0 ± 0.5 |
| **NATA2** | 100 µM acCoA + Jasmonate isoleucine | 64.5 ± 0.8 |
| **NATA2** | 100 µM acCoA + ***γ***-aminobutyric acid | 67.5 ± 0.5 |
| **NATA2** | 100 µM acCoA + Glutamate | 67.5 ± 0.3 |

**Supplementary Table 3A/B. Thermal melting point (*Tm*) of NATA1Δ and NATA2Δ established by DSF.** Analysis was carried out in buffer Tris8.5.

**A**

| Target Gene | Purpose | | Primer sequence |
|---|---|---|---|
| *NATA2* (At2g39020) | Cloning | forward | GGG GAC AAG TTT GTA CAA AAA AGC AGG CTT CAT GGC AGC CGC CGC ACC G |
| | Cloning | reverse | GGG GAC CAC TTT GTA CAA GAA AGC TGG GTC CTA GAT GTT GAC CTG ATC AAA AGC TTC |
| SALK-LBa1 | Confirm T-DNA insertion | SALK primer | TGGTTCACGTAGTGGGCCATC G |
| *NATA2* (At2g39020) | Confirm T-DNA insertion | T-DNA forward | TCA ACT CAC ACA CAT CGT GTG |
| *NATA2* (At2g39020) | Confirm T-DNA insertion | T-DNA reverse | AAC AAC CCA TTC CAC TCT TCC |
| *NATA2*-forward | Confirm NHEJ mutagenesis | forward | TGG GTT TGT TCT GTT TTT C |
| *NATA2*-reverse | Confirm NHEJ mutagenesis | reverse | TCTCAGCAACACAGCATCA A |
| Hyg resistant gene | Confirm TALEN | forward | GCT CCA TAC AAG CCA ACC AC |
| | Confirm TALEN | reverse | CGA AAA GTT CGA CAG CGT CTC |

**B**

| Target Gene | | Primer sequence |
|---|---|---|
| *HSP70.1* (At5g02500) | forward | GGGGAAGATTTTGACAACA |
| | reverse | TCTTCGCTCTCTCACAGGAAG |
| *PR1* (At2g14610) | forward | TGATCCTCGTGGGAATTATGT |
| | reverse | TGCATGATCACATCATTACTTCAT |
| *PR2* (At3g57260) | forward | AGCTTCCTTCTTCAACCACACAGC |
| | reverse | TGGCAAGGTATCGCCTAGCATC |
| *PR5* (At1G75040) | forward | AGCAATGCCGCTTGTGATGAAC |
| | reverse | ATCACCCACAGCACAGAGACAC |
| *NATA2* (At2g39020) | forward | CGATTGATGATCCGGAGAGT |
| | reverse | AGCGGTGAAGATGGGTTATG |
| *NATA1* (AT2G39030) | forward | AGCAGATGGGTGCGCAGGTT |
| | reverse | TCGCTCGATGGGTCTCATGCA |
| *ACT2* (At3g18780) | forward | TCCCTCAGCACATTCCAGCAGAT |
| | reverse | AACGATTCCTGGACCTGCCTCATC |

**Supplementary Table 4A/B. Primers used A)** Primers used for cloning and genotyping. **B)** Primers used for quantitative RT-PCR.
